# Regulated control of gene therapies with a drug induced switch

**DOI:** 10.1101/2020.02.21.956664

**Authors:** Alex Mas Monteys, Amiel A Hundley, Paul T Ranum, Euyn Lim, Luis Tecedor, Amy Muehlmatt, Beverly L Davidson

## Abstract

To date, gene therapies for human application rely on engineered promoters that cannot be finely controlled. Here, we report a universal switch element that allows precise control for gene silencing or gene replacement after exposure to a small molecule. Importantly, these small molecule inducers are in human use, are orally bioavailable when given to animals or humans, and can reach both peripheral tissues and the brain. Moreover, the switch system (X^on^) does not require the co-expression of any regulatory proteins. Using X^on^, translation of desired elements for gene knockdown or gene replacement occurs after a single oral dose, and expression levels can be controlled by drug dose or in waves with repeat drug intake. This universal switch can provide temporal control of gene editing machinery and gene addition cassettes that can be adapted to cell biology applications and animal studies. Additionally, due to the oral bioavailability and safety of the drugs employed, the X^on^ switch provides an unprecedented opportunity to refine gene therapies for more appropriate human application.

## Introduction

While viral and nonviral approaches for gene therapies have made tremendous advancements over the last twenty years, the major focus has been on the cargo delivery system; *e.g*., viral capsid evolution and engineering for adenoassociated viruses (AAVs), expanding the landscape of cell-targeting envelopes for lentiviruses, and refining lipid nanoparticles for improved uptake. However, the cargo itself, and more importantly the elements controlling the expression from that cargo, have been largely untouched aside from using engineered promoters or 3’ regulatory elements to restrict expression to certain cell types^1,2^.

To address this gap, we developed X^on^, a method to finely control protein expression via a drug inducible switch. Importantly, X^on^ does not require any bacterial or other external elements for regulation. The X^on^ system can be applied to any genetic element of interest in cells or animals, and takes advantage of drugs that are orally bioavailable and in human use.

## Results

Initial experiments to test our approach were done using the actual target of the drugs, the *SMN2* gene, for Spinal Muscular Atrophy (SMA) therapy. SMA is due to mutations in the gene *SMN1*. Humans have a very similar gene, *SMN2* that can serve to modify the severity of SMN1 deficiency depending on the number of *SMN2* copies resident in the patient’s genome. *SMN2* cannot fully replace *SMN1*, however, because unlike *SMN1*, *SMN2* has undergone variations impairing exon7 inclusion. As a consequence, only ~10% of SMN2 is correctly spliced^3,4^. Two drugs, LMI070^5^ and RG7800/RG7619^6^, can improve exon7 inclusion and are in later stage clinical testing. We reasoned that the exon6/7/8 cassette could be co-opted and refined to control the expression of a gene of interest, rather than SMN2 protein.

For this, SMN2-on cassettes were generated to provide drug-induced reporter gene expression. HEK293 cells were transfected with the SMN2-on expression cassettes and luciferase activity evaluated in response to varying doses of LMI070 or RG7800. As depicted in Fig 1a, firefly luciferase activity would be expected with exon 7 inclusion, while exon 7 exclusion results in the presence of a premature stop codon and lack of signal. The SMN2-on switch was tested in its native format, or altered for constitutive inclusion or reduced background exon 7 inclusion by modifying SMN2 exon 7 donor or acceptor splice sites, respectively (Fig. 1b-c). Both RG7800 and LMI070 induced luciferase activity from the SMN2-on minigene, with a complete splicing switch evident at drug concentrations greater than 1 μM (Fig. 1c, Extended Data Fig. 1a,b) and an overall induction of approximately 20-fold for the refined SMN2-on switch system (Fig 1d,e, Extended Data Fig. 1c). In the setting of the SMN2-on cassette, LMI070 was more active than RG7800 (Fig 1d, Extended Data Fig. 1a,b), which may be due to their different mechanisms of action^7,8^.

**Fig 1.**
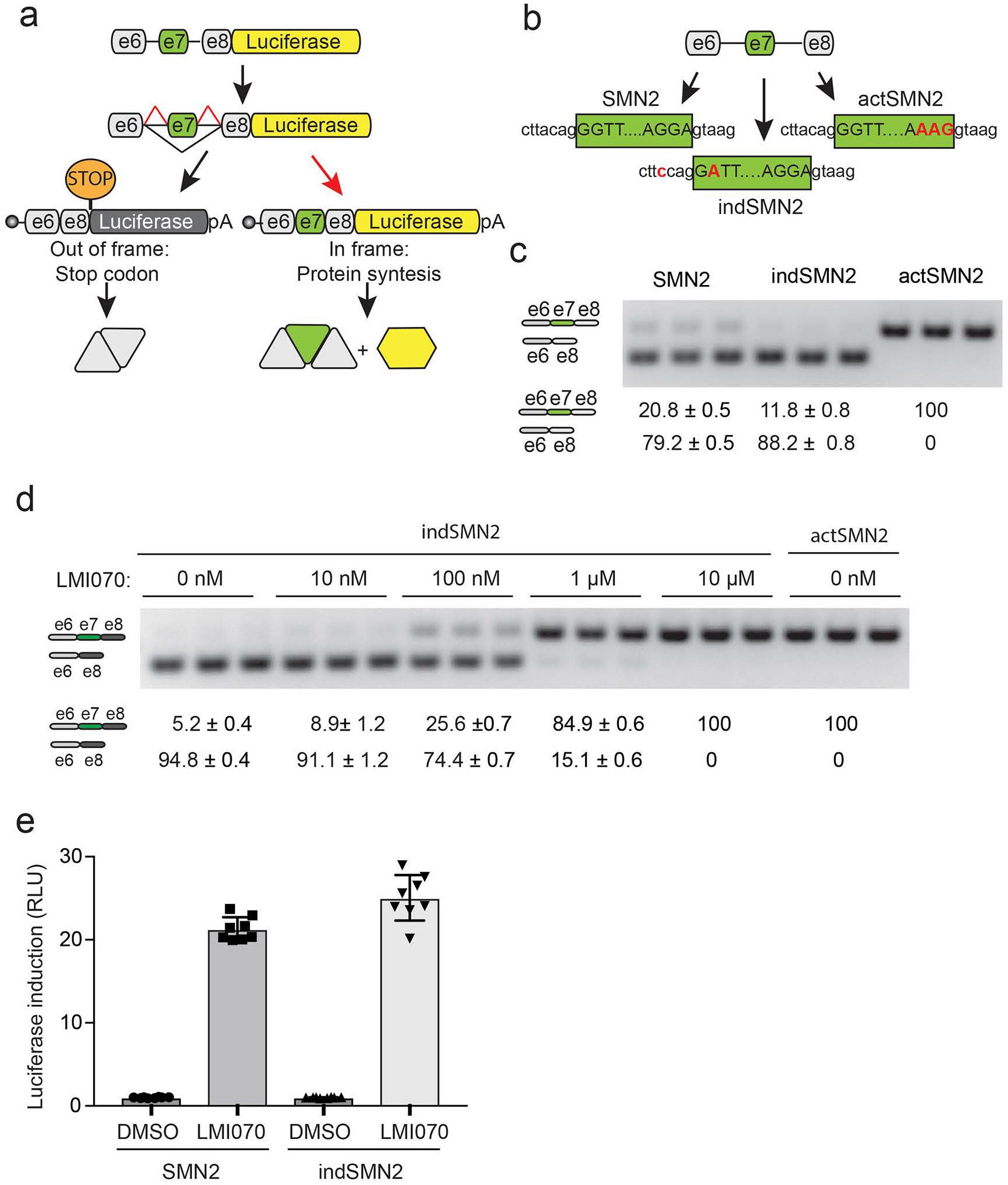
Generation and assessment of the SMN2-on cassette. a) Cartoon depicting the SMN2-on cassette and its mechanism of action. In the absence of exon 7 (e7), a premature stop codon in the exon 6/8 transcript (e6/8) blocks translation of the luciferase cDNA sequence, while inclusion of the alternative splice exon 7 by LMI070 or RG7800 (depicted by red arrow) permits translation of the e6/7/8:*Firefly* luciferase fusion protein. b) cartoon depicting SMN2 exon 7 in its native sequence, or with the splice site modifications introduced for constitutive inclusion (5’ donor splice site, actSMN2) or for reduced background levels of exon 7 inclusion (3’ acceptor splice site, indSMN2). c) Representative RT-PCR reaction showing exon 7 inclusion with the SMN2-on cassettes in the absence of LMI070. The quantification of the exon 7 spliced-in or spliced-out transcripts are depicted, and are the mean ± SEM of 6 biological replicates. d) Representative RT-PCR reaction showing exon7 inclusion of the indSMN2 cassette in response to different concentrations of LMI070. The quantification of exon 7 spliced-in or spliced-out transcripts are the relative transcript levels (mean ± SEM of 9 biological replicates. e) *Firefly* luciferase induction of the SMN2 and indSMN2 cassettes in response to LMI070 (100 nM) relative to control DMSO treated cells. The fold change of luciferase activity in LMI070-treated samples relative to DMSO treated control cells is shown. All samples are normalized to *Renilla* luciferase activity. The data are the mean ± SEM of 8 biological replicates.

We next set out to test for exons that were more sensitive to LMI070 to reduce non-target splice events. We treated HEK293 cells with 25 nM LMI070 for 12 hours and ascertained the splicing changes induced using RNA-Seq. Reads obtained from 4 control and 4 LMI070 treated samples were aligned to the genome using the STAR aligner^9^ and splicing events exclusive to the LMI070 treated samples identified (Fig. 2a, Extended Data Table 1, Extended Data Fig. 2 a-h). A total of 45 novel splicing events were identified following LMI070 treatment that were above our threshold of an average of greater than 5 novel intron splicing events in LMI070 treated samples (Fig. 1, Extended Data Table 1). Among them, 23 events were found exclusively and in all LMI070 treated samples, and the remaining 22 were evident in all treated and in one of the control untreated samples (Extended Data Table 1). To assess exclusivity of the 45 identified candidate positions to LMI070 treatment the chromosomal locations were assessed in Intropolis^10^, a resource containing a list of all exon-exon junctions found in 21,504 human RNA-Seq datasets. In one example, the canonical exon-exon junction in *SF3B3* (Fig. 2b; Extended Data Table 1) was observed in 12,872 datasets at an average frequency of 64 counts per dataset, while the LMI070 induced splice event was observed in 10 and 1 dataset(s), respectively, for the 5’ and 3’ exon-exon junctions. The average counts per dataset for the 5’ exon-exon junction was 1.3 while the 3’ exon-exon junction was observed only once in all 21,504 sets (Extended Data Table 1).

**Fig 2.**
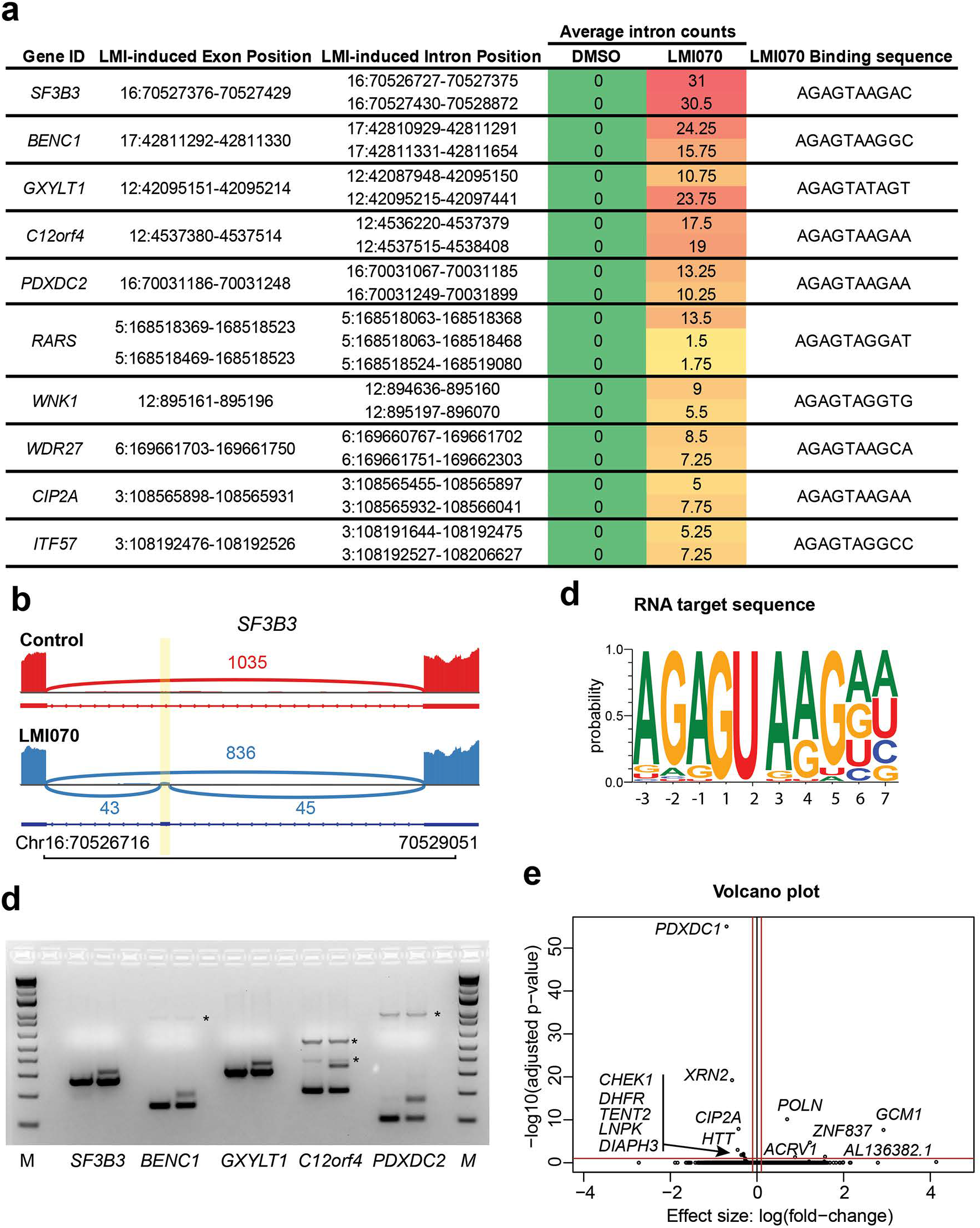
RNA-Seq for LMI070-responsive pseudo exon discovery. a) Table showing the top candidate spliced in events identified by RNA-Seq in HEK293 cells treated with 25 nM LMI070. Shown are Gene ID, the LMI070-induced exon and the flanking intron positions, average intron counts, and the U1 LMI070-targeted binding sequence. b) Sashimi plot depicting the SF3B3 splicing event in the absence (red) or in response (blue) to 25 nM LMI070. The genomic location of the LMI070 spliced-in exon for SF3B3 is indicated (yellow bar), and the number of intron counts indicated. c) Sequence logo of the U1 RNA binding sequence targeted by LMI070 from 45 spliced-in exons identified by RNA-Seq. d) cDNAs amplified from HEK293 cells treated with DMSO or LMI070 shows spliced-in events for *SF3B3*, *BENC1*, *GXYLT1*, *C12orf4* and *PDXDC2*, which were confirmed by Sanger sequencing. Asterisks mark nonspecific bands amplified from the PCR reaction. e) Volcano plot illustrating the differentially expressed genes between DMSO- and LMI070-treated cells. The horizontal bar extending from the y axis represents the significance, 0.05, plotted on a -log10 scale. Thresholds of −0.1 and 0.1-fold-change are indicated by red vertical bars. Genes that meet the threshold for significance and minimum fold change requirements are labeled.

The pseudoexons identified in LMI070 treated samples share a strong 3’ AGAGUA motif consistent with the previously identified U1 RNA binding site targeted by LMI070 (Fig. 2c)^5^. To experimentally validate the identified LMI070 induced splicing events, primer pairs binding the flanking exons of the top 5 candidate genes (Fig.2a, Extended Data Fig. 2a-e) were generated. Robust amplification of the novel exons was detected by PCR exclusively on cDNA samples generated from HEK293 cells treated with LMI070 (Fig. 2d). We evaluated the impact of LMI070 treatment on global gene expression by assessing differential expression analysis. DESeq2 revealed strikingly few differentially expressed genes with only 6 upregulated and 24 downregulated genes passing our threshold for significance (p< 0.05; Benjamin-Hochberg multiple testing correction)^11^. After filtering out genes with low fold-change values (<0.1 fold) only 5 upregulated and 9 downregulated genes were identified (Fig. 2e).

We next developed a series of switch-on cassettes from the top 4 LMI070-responsive exons found in our RNA-Seq dataset. For this, the minimal intronic intervening sequences necessary to recapitulate splicing of pseudoexons in *SF3B3*, *BENC1*, *C12orf4* and *PDXDC2* were cloned upstream of luciferase or eGFP cDNAs (Fig. 3a,c). To limit translation only in response to the drug, a Kozak sequence followed by an AUG start codon were positioned within the novel exon to be included in response to LMI070 binding (Fig. 3a,c). HEK293 cells were transfected with the candidate cassettes, treated with LMI070, and luciferase activity or eGFP expression determined 24h later. Increased luciferase expression was observed for each candidate cassette in response to LMI070, with the SF3B3-on switch showing a more than 100-fold induction (Fig 3b). Notably, this is 5x the 20-fold induction afforded by the SMN2-on cassette (Fig 1e). Similarly, eGFP expression was only detected in cells transfected with the SF3B3-on-eGFP cassette in response to LMI070 treatment (Fig 3d).

**Fig 3.**
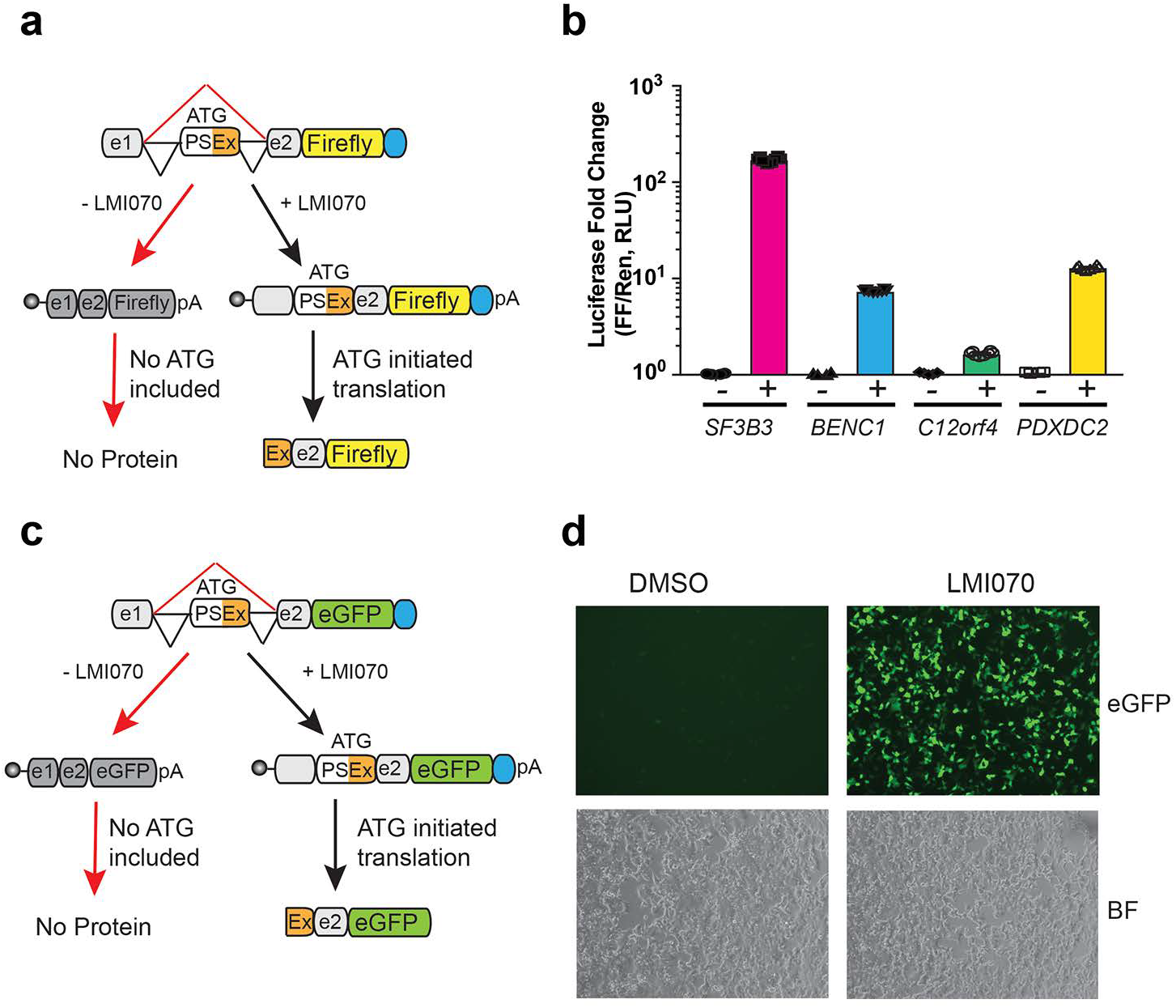
Candidate minigene cassette-responses to LMI070. a) Cartoon depicting the candidate minigene configuration for controlling translation of *Firefly* luciferase, with e1 referring in all cases to the exon 5’ of the LMI070-induced pseudoexon from the RNA-Seq analysis (see Fig. 2b, Extended Data Fig 2a-i). A Kozak and ATG initiation codon were positioned within the LMI070-spliced-in exon to initiate translation only in response to drug. b) Luciferase induction of the minigene cassettes for *SF3B3*, *BENC1*, *C12ORF4* and *PDXDC2*. The fold change luciferase activity in LMI070-treated samples (depicted as +) is relative to DMSO-treated (depicted as -) cells, with data normalized to *Renilla* luciferase. Data are the mean ± SEM of 8 biological replicates. c) Cartoon depicting the minigene cassette controlling eGFP expression. A Kozak and ATG initiation codon were positioned within the LMI070-induced exon to initiate translation only after treatment with drug. d) eGFP expression in HEK293 cells transfected with the SF3B3 minigene cassette (X^on^-eGFP) and treated 24 hr later with DMSO (left) or LMI070 (right).

While there was measurable baseline luciferase activity in the absence of LMI070 for all candidate cassettes, the SF3B3 cassette had the least background (Extended Data Fig 3a). As recent work using ribosome foot printing revealed that non-AUG start codons provide for translation initiation^12^, we evaluated the presence of in frame non-AUG start codons that could drive luciferase translation in the absence of drug. We found that all cassettes contained in frame non-AUG codons, however there was only one in frame non-AUG codon in the SF3B3 cassette (Extended Data Fig 3b).

To assess the baseline splicing from the engineered switch cassettes two different PCR assays were done, one with primers binding the flanking exons to detect all transcripts, and another with a primer binding within the novel spliced exon (Extended Data Fig 3c). The LMI070-spliced exon was not detected with primer pairs binding the flanking exons, whereas there was faint signal when priming specifically the novel exon sequence (Extended Data Fig 3c). Overall, our results suggest that in the absence of LMI070 the alternative exon may be included in a small fraction of the transcripts, mirroring what was found in the Intropolis dataset (Extended Data Table 1).

Splicing machinery selection of 5’ and 3’ splice sites pairs is defined by several cis-acting sequences that collectively comprise the ‘splicing code’, including combinations of silencer and enhancer splicing sequences that repress or promote the selection of cryptic or correct splice sites. Using the Human Splicing Finder website^13^, we screened the *SF3B3* intron sequence for putative silencer and enhancer sequences that could modulate inclusion of the SF3B3 LMI070-spliced pseudoexon (Extended Data Fig 3d). Three intronic regions rich in silencing sequences that could repress splicing in the absence of the drug were identified downstream of the pseudoexon. To test their impact on drug-induced control, SF3B3-on-reporter cassettes containing the full intron sequence (SF3B3int), intron fragments rich in silencer sequences (SF3B3i1, SF3B3i2, and SF3B3i3), or an intron fragment less enriched for intronic silencer sequences (SF3B3i4) were generated (Extended Data Fig 3e). HEK293 cells were transfected with the original SF3B3-on switch or the cassettes containing alternate intronic sequences and splicing and luciferase activity determined 24h later. Whereas cells transfected with SF3B3i4 showed activity similar to the original SF3B3-on construct, luciferase activity was reduced in all the other groups irrespective of drug treatment (Extended Data Fig 3f,g). The fold-change in luciferase activity in response to LMI070 was significantly higher in cells transfected with plasmids containing the original intron (>200 fold, Extended Data Fig 3g). Surprisingly, the reduction of the luciferase activity was not related to splicing repression of the novel exon, but to the generation of additional spliced transcripts in the absence of the drug (Extended Data Fig h,i). The original SF3B3-on cassette (hereafter referred to as X^on^) was used for all further studies.

Next, we tested X^on^ for responsiveness when expressed from promoters of varying strengths. HEK293 cells were transfected with expression plasmids containing the X^on^ luciferase encoding cassettes under the control of the Rous sarcoma virus (RSV), the phosphoglycerate kinase (PGK), or the minimal cytomegalovirus (mCMV) promoter. All promoters drove inducible expression (Fig. 4a), with a clear dose response in fold-change luciferase activity (Fig. 4b, Extended Data 4a,b) that was mirrored by splicing assessment (Fig. 4c, Extended Data 4c). Overall these constructs provide a gradient of induction with RSV>PGK>mCMV.

**Fig 4.**
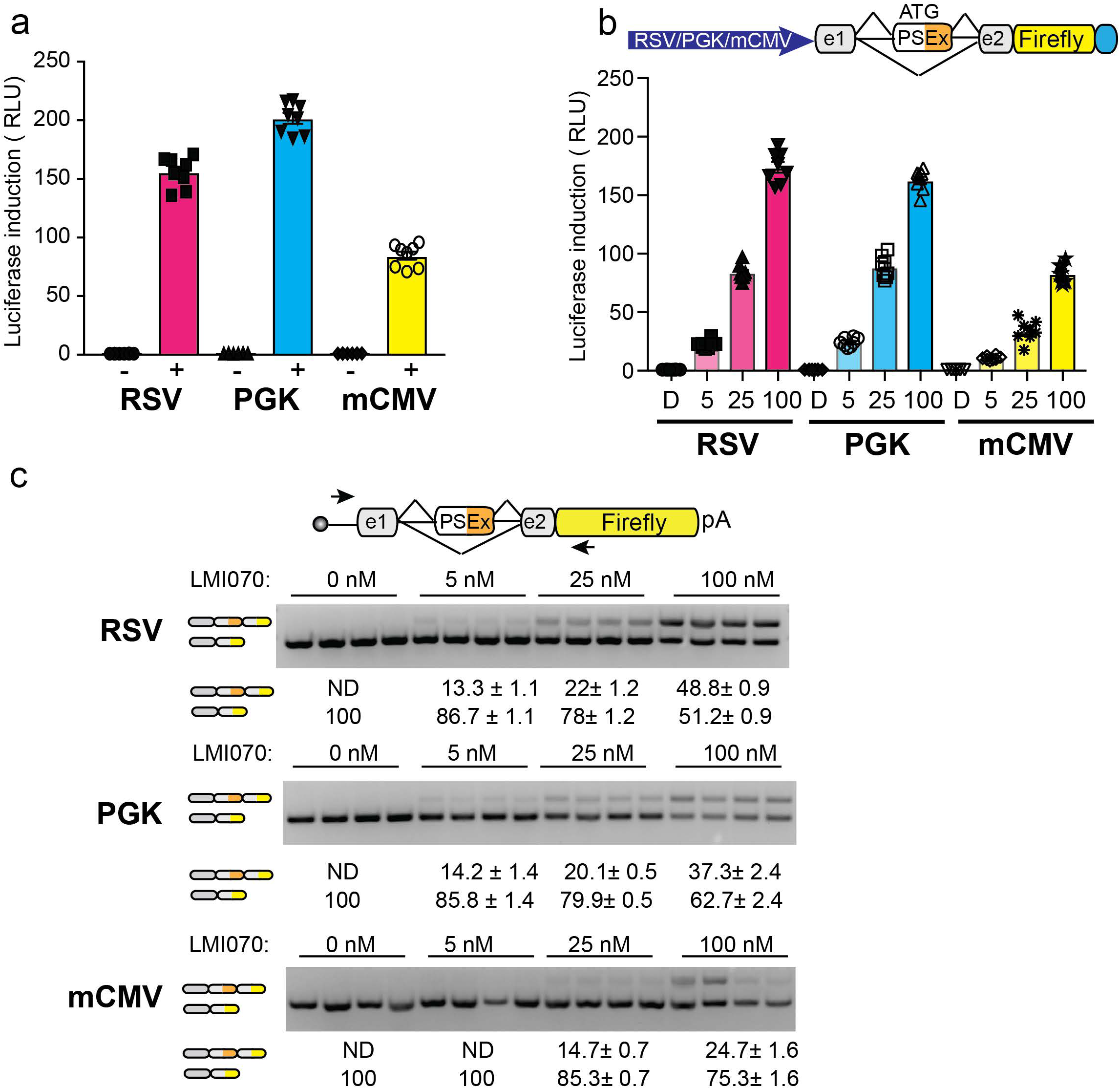
Activity of the SF3B3-X^on^ cassette when expressed various promoters and their responsiveness to LMI070 dose. a) Luciferase induction after transfection of plasmids containing the noted SF3B3-X^on^ -luciferase expression cassettes into HEK293 cells followed by treatment with LMI070 (100 nM, denoted as plus) or treated with DMSO (denoted as minus). All samples are normalized to *Renilla* luciferase activity and are relative to DMSO treated cells. Data are the mean ± SEM of 8 biological replicates. b) Luciferase induction in HEK293 cells transfected with plasmids containing the noted SF3B3-X^on^ cassettes and treated with varying doses of LMI070. All samples are normalized to *Renilla* luciferase activity and are relative to DMSO treated cells (0 nM). Data are the mean ± SEM of 8 biological replicates. c) Representative gels from RT-PCR analysis for assessment of the LMI070-induced pseudoexons expressed from the noted promoters in response varying doses of LMI070. Pseudoexon inclusion was detected using primers flanking the pseudoexon. Splicing was quantified and transcript levels presented as the mean ± SEM of 8 biological replicates.

To assess the X^on^ system *in vivo*, an AAV X^on^ vector was developed and packaged into AAV9 (AAV9.RSV.X^on^.eGFP). AAV9.X^on^.eGFP (3E11 vg/mouse) was administrated intravenously (IV) to mice, and 4 weeks later animals were given a single dose of vehicle or LMI070 at 5 or 50 mg/kg and eGFP expression assessed 24 h later (Fig. 5a). There was notable eGFP expression in sections from liver (Fig. 5b). The dose response in eGFP signal noted in tissues by microscopy was confirmed by western blot (Fig. 5c, Extended Data Fig. 5a) and splicing assay (Fig. 5d). Cumulatively, these data show that *in vivo*, the X^on^ cassette can be used to drive gene expression from AAV vectors after a single administration of drug. Importantly, protein levels and novel exon splicing directly correlated to the LMI070 dose.

**Fig 5.**
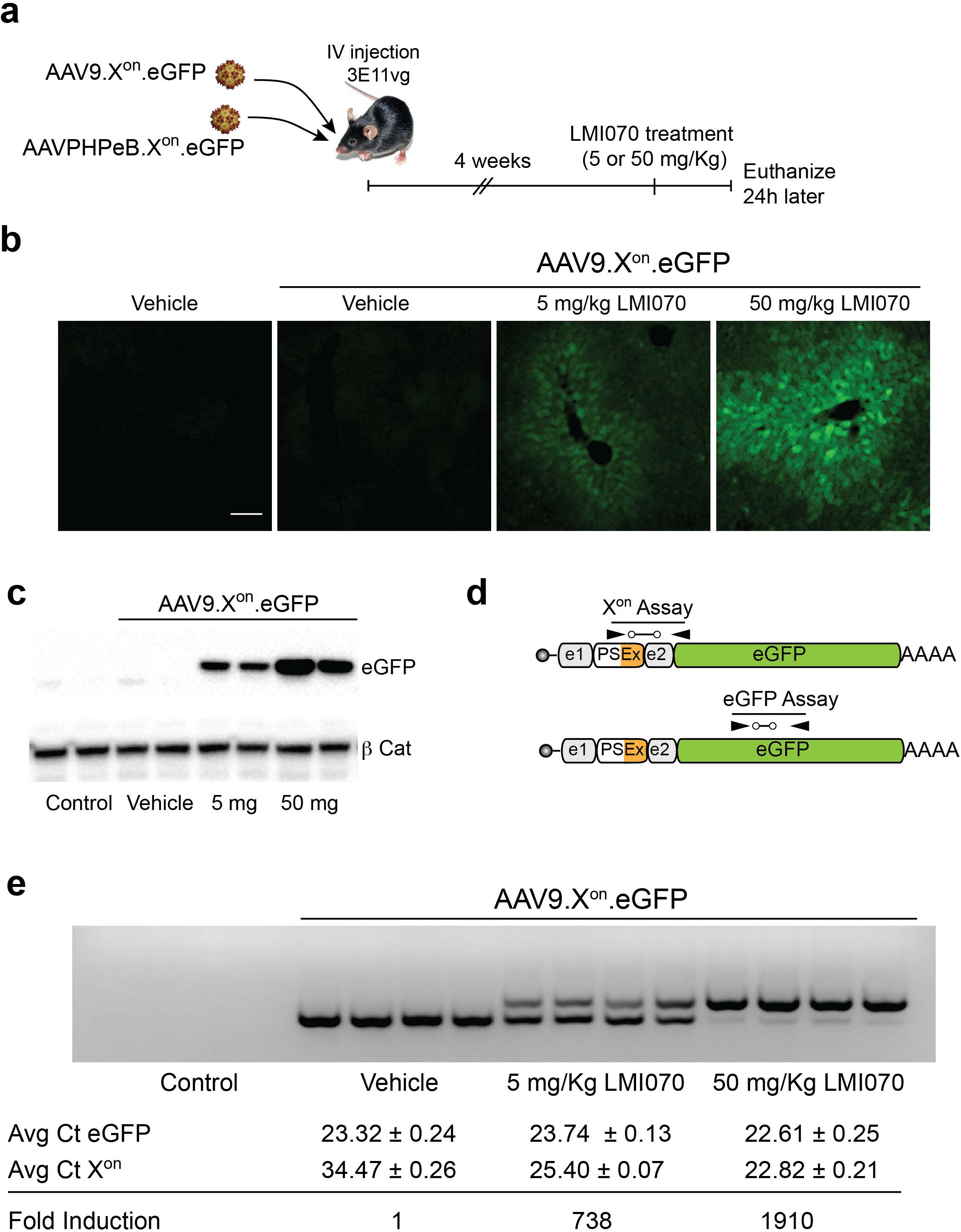
*In vivo* activity of X^on^. a) Schematic of the *in vivo* studies using either AAV9.X^on^.eGFP or AAVPHPeB.X^on^.eGFP. Mice were injected iv and 4 weeks later were treated with a single dose of 5 or 50 mg/kg LMI070, and tissues harvested 24h later to assess splicing, transcript levels and protein expression b) Representative photomicrograph of liver tissue sections showing eGFP in liver 24h after treatment with LMI070 at 5 or 50 mg/Kg (scale bar 100 µm). c) Representative western analysis of eGFP protein levels (2 samples shown; 4 mice/group). β-catenin is shown as loading control. d) Cartoon depicting the X^on^ assays designed to quantify the LMI070-induced transcripts and eGFP expression levels from the X^on^ cassette after AAV9.X^on^.eGFP gene transfer. e). Representative gel from PCR assays demonstrates inclusion of the splicing activity in response LMI070. Data shows the average Ct values for eGFP or LMI070-induced expression using the X^on^ gene expression assays depicted in d). Fold change of the spliced expression cassette is shown relative to basal levels in mice injected with AAV9.X^on^.eGFP and treated with vehicle.

To assess the applicability of our system for brain targeted gene therapies, the X^on^ was packaged into AAVPHPeB^14^ (AAVPHPeB.X^on^.eGFP) and delivered IV to mice (3E11 vg/mouse). Again, eGFP expression and novel exon splicing were evident only in response to drug, and in dose responsive manner (Extended Data Fig 5b,c).

While gene editing approaches provide an enormous opportunity for altering or removing disease alleles, prolonged expression of the editing machinery from viral vectors could be problematic. The editing enzymes are foreign proteins and may induce immune responses, and prolonged gene expression would increase opportunities for off-target editing^15,16^. We tested the utility of the X^on^ system to regulate editing, using *huntingtin* (*HTT*), a target for gene silencing approaches for Huntington’s disease (HD)^17^, as an example.

First, we designed gRNAs to mediate SaCas9-mediated deletion of mutant *HTT* (m*HTT*) exon 1. For this, we took advantage of a single nucleotide polymorphism (SNP) 5’ to m*HTT* exon 1 that creates a SaCas9 protospacer adjacent motif (PAM; sg935, Fig. 6a, Extended Data 6a). When used in combination with a sgRNA targeting the downstream intron (sgi3, Fig. 6a, Extended Data 6a)^18^, these gRNAs edit *HTT* exon1 via SaCas9 and reduce HTT mRNA and protein levels (Extended Data Fig. 6b-e). Next, the X^on^ switch for drug-inducible SaCas9 expression was generated, and compared to the constitutively active cassette (Fig. 6b). There was a clear dose response of SaCas9 expression with LMI070 (Fig. 6c). X^on^-SaCas9 plus the relevant gRNAs were then transfected in HEK293 cells and the *HTT* locus, HTT mRNA and HTT protein levels assessed (Fig. 6d). There was equivalent editing between the samples treated with LMI070 and the samples constitutively expressing SaCas9. (Fig. 6e). More importantly, there was a concomitant reduction at the RNA level upon LMI070 treatment, with *HTT* transcripts reduced by 50%, similar to the extent noted in cells transfected with the constitutively active editing expression cassette (Fig. 6f). Protein levels were similarly reduced (Fig. 6g). And while minimal editing was detected on cells transfected with active gRNAs and X^on^-SaCas9 treated with DMSO (Fig. 1e), transcript and protein levels remained unaffected (Fig. 6f,g). Together, our data show that the X^on^ switch together with allele specific gRNAs directed to m*HTT* provides an important advance to HD treatment.

**Fig 6.**
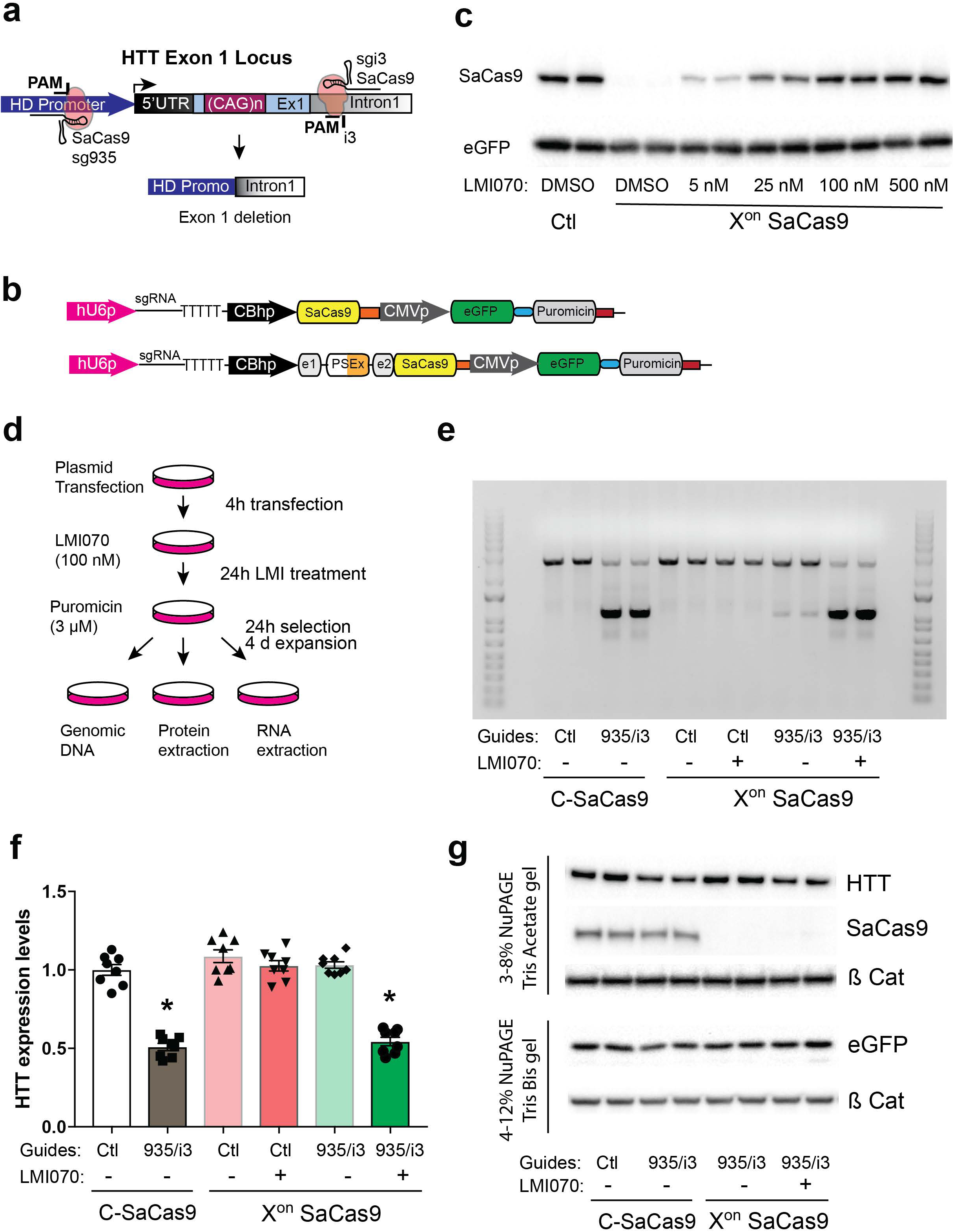
Regulated SaCas9 editing for Huntington’s disease. a) Cartoon depicting the allele-specific editing strategy to abrogate mutant HTT expression. SNPs within PAM sequences upstream of HTT exon-1 permit targeted deletions of the mutant allele when present in heterozygosity. After DNA repair, mutant HTT exon-1 is deleted by a pair of sgRNA/Cas9 complexes binding upstream and downstream of exon 1. b) Cartoon depicting the plasmids used to co-express the sgRNA sequences, SaCas9 (constitutively or with the X^on^ switch), and the selective reporter eGFP/puromycin expression cassettes. c) Bicistronic plasmids containing the CBh.X^on^.SaCas9, or the CBhpSaCas9 expression cassette together with CMVeGFP expression cassettes were transfected in HEK293 cells and treated with varying doses of LMI070 and SaCas9 levels determined by western blot 24h later. eGFP protein levels served as transfection control. Blot is representative of 2 of 4 technical replicates. d) Schematic of regulated CRISPR for assessing *HTT* editing. HEK293 cells were transfected with plasmids expressing the sgRNA sequences, SaCas9 (constitutively or via the X^on^ switch), and the selective reporter eGFP/puromycin expression cassettes. Four hours later cells were treated with 100 nM LMI070, and 24 later with 3 µM Puromycin. After 24 hours of selection cells the Puromycin was removed and cells expanded for 4 days after which cells were collected for genomic DNA, protein and RNA isolation. e) Representative gel for 2 of 5 biological replicates) depicting *HTT* exon 1-targeted deletion assessed by PCR assay after transfection with constitutive or X^on^-SaCas9 CRISPR plasmids using control (Ctl) sgRNAs or 935/i3 sgRNA sequences and the selective reporter eGFP/puromycin expression cassettes. f) Transcript levels as assessed RT-qPCR analysis of HTT mRNA levels from HEK293 cells transfected with constitutive or X^on^-SaCas9 CRISPR cassettes. Samples are normalized to human GAPDH, and data are the mean ± SEM relative to cells transfected with plasmids containing the constitutive SaCas9 CRISPR cassette expressing a control sgRNA sequence (8 biological replicates; *p < 0.0001, one-way ANOVA followed by a Bonferroni’s post hoc). g) Representative western blot (2 of 4 biological replicates) for HTT, SaCas9 and eGFP protein levels after puromycin selection and expansion of HEK293 cells transfected with constitutive or X^on^-SaCas9 CRISPR cassettes. Cells transfected with constitutive cassettes containing sgRNA expression cassette were used as a control. β-catenin served as loading control.

In summary, we present a simple, highly adaptable tool for regulated gene expression for *in vitro* cell biology applications and *in vivo* evaluation of any protein and testing of new therapies. We show the utility of the X^on^ system for gene addition and gene editing, and demonstrate its exquisite control in cells in culture, or in tissues via AAV delivery. Researchers can use X^on^ in cells or animals to test for additional waves of expression, and use different promoters and drug doses for varying levels of inducibility. The X^on^ tool also gives researchers the means to test gene products that when constitutively expressed are toxic. Indeed, the X^on^ system can be applied to any biological question where fine expression control is desired.

## Methods

### Cell culture, transfection and LMI070/RG7800 treatment

Human embryonic kidney (HEK293) cells (obtained from CHOP Research Vector Core stock) were maintained in DMEM media containing 10% Fetal Bovine Serum (FBS), 1% L-Glutamine and 1% penicillin/streptomycin at 37 °C with 5% CO_2_. Cells were cultured in 24 well plates and transfected at 80-90% confluence using Lipofectamine 2000 transfection reagent, according to the manufacturer’s protocol. For all experiments, 4 hours after plasmid transfection, cells were treated with LMI070 (MedChemExpress, HY-19620, suspended in DMSO) or RG7800 (MedChemExpress, HY-101792A, suspended in H_2_O) at the indicated concentrations. Cells were tested for mycoplasma by Research Vector Core. None of the cells used in the study were listed in ICLAC database of commonly misidentified cell lines.

### Plasmids, primers and custom made TaqMan gene expression assays

All plasmids, primer sequences, and custom Taqman gene expression assays to determine SF3B3 novel exon inclusion are available upon request. Primers and custom Taqman gene expression assays were obtained from IDT Integrated DNA Technologies.

### *In vitro* luciferase assays

HEK293 cells were cultured in DMEM (10% FBS (v/v), 1% Pen/ Strep (v/v), and 1% L-glutamine (v/v) in a 24-well plate. At 70%–80% confluence, cells were co-transfected with the X^on^. *Firefly* luciferase cassettes (0.3µg/well) and a SV40p-*Renilla* luciferase cassette as transfection control (0.02µg/well). Four hours after transfections cells were treated either with LMI070 or RG7800 at indicated concentrations. At 24 h after transfection, cells were rinsed with ice-cold PBS and *Renilla* and *Firefly* luciferase activities were assessed using the Dual-Luciferase Reporter Assay System (Promega) according to the manufacturer’s instructions. Luminescent readouts were obtained with a Monolight 3010 luminometer (Pharmigen). Relative light units (RLUs) were calculated as the quotient of *Renilla/Firefly* RLUs and results expressed relative to mock treated control cells.

### Animals and Histology

Animal protocols were approved by The Children’s Hospital of Philadelphia Institutional Animal Care and Use Committee. Five to six-week-old male C57Bl6/j mice were obtained from Jackson Laboratories (Bar Harbor, ME, USA). AAV vectors (AAV9. X^on^.eGFP or AAVPHBeB.X^on^.eGFP; generated at the CHOP Research Vector Core) were administrated by retroorbital injection at 7-8 weeks of age with a total of 3E11 vg in150 μl infused. After 4 weeks, a single dose of LMI070 (5 or 50mg/kg, MedChemExpress HY-19620) or vehicle solution was administrated by oral gavage. After 18-24h, mice used for biochemical or molecular studies were anesthetized and perfused with 0.9% cold saline mixed with 2-ml RNAlater (Ambion) solution. Brains and liver samples were collected, flash frozen in liquid nitrogen, and stored at −80 C until use. For immunohistochemistry studies, mice were perfused with 15 mL ice-cold 0.1M PBS followed by 15 mL 4% paraformaldehyde. For brain sections, eGFP visualization was done by IHC using rabbit anti-GFP antibody (Invitrogen, 1:200) followed by Alexa488-conjugated goat anti-rabbit (Invitrogen, 1:500) and Alexa488-conjugated chicken anti-goat (Invitrogen, 1:500). Slices were mounted on Superfrost plus slides and coverslips mounted using fluoro-gel mounting media. Sections were analyzed using a DM6000B Leica microscope equipped with a L5 ET filter cube (ex:em of 470±20:525±15 nm and dichroic 495 nm), a 20X HC APO PLAN (N.A. 0.70) lens connected to a Sola Light Engine LED light source (Lumencor). Images were collected with a Hammatsu Orca flash4.0 monochrome camera controlled by Leica LAS X (v.3.0.3) software. Brain images represent a 7.98 µm thick z-stack deconvoluted by 3 iterations of the Blind algorithm.

### RNA extraction, RT-PCR and Splicing assays

Total RNA was extracted using Trizol (Life Technologies) according to the manufacturer’s protocol, with the exception of 1 µL Glycoblue (Life Technologies) in addition to the aqueous phase on the isopropanol precipitation step and a single wash with cold 70% ethanol. To determine HTT expression levels after transfection, RNA samples were quantified by spectrophotometry and subsequently cDNAs generated from 1 mg of total RNA with random hexamers (TaqMan RT reagents, Applied Biosystems). To determine human HTT expression levels in HEK293 cells, we used TaqMan probes for human HTT and glyceraldehyde 3-phosphate dehydrogenase (GAPDH) mRNAs obtained from Applied Biosystems. Relative HTT gene expression was determined using the ddCt method. To determine splicing of the SMN2-on and X^on^ switches, 2 µg of total RNA from HEK293 cells or tissue samples was treated with DNAseI Free kit (Thermofisher) followed by cDNA generation using the High capacity cDNA kit (Thermofisher). Splicing was determined by PCR using the Phusion High-Fidelity polymerase (Thermofisher) and PCR products separated on a 2.5% agarose gel pre-stained with EtBr and spliced-in and spliced-out band densitometry performed using with the ChemiDoc Imaging System (BioRad) and Image Lab analysis software. Splicing induction from mouse tissues was determined using two custom TaqMan assays designed to determine total or LMI070-spliced in mRNA transcripts. The percentage of induction was determined by dividing the average Ct novel exon and average Ct Total, relative to control animals injected with AAV virus plus vehicle.

### Western Blots

HEK293 cells were rinsed and lysed with Passive Lysis Buffer (PBL, Promega), protein concentrations determined using the DC protein assay (Bio-Rad), and 10-20 µg of protein loaded on a 3%–8% NuPAGE Tris-Acetate Gel in tris-acetate buffer (Novex Life Technologies) or a 4-12% NuPAGE TrisBis NuPAGE gels in MES buffer (Novex Life Technologies) to determine HTT, SaCas9, or eGFP protein levels, respectively, with β-catenin as loading control. Livers were homogenized in RIPA buffer (final concentration: 50 mM Tris, 150 mM NaCl, 1% Triton-X100, 0.1% SDS, 0.5% sodium deoxycholate, with Complete protease inhibitors (Roche) and samples incubated for 1 hr rotating at 4°C then clarified by centrifugation at 10,000 x g for 10 minutes. Total protein concentration was determined by DC protein assay (BioRad) and 30 µg loaded on a 4-12% NuPAGE Bis-tris gels in MES buffer (Novex Life Technologies) to determine eGFP and β-catenin levels. After electrophoresis, proteins were transferred to 0.2 µm PVDF (Bio-Rad). Membranes were blocked with 5% milk in PBS-T and then blotted with a mouse anti-HTT (MAB2166. dilution: 1:5,000; Millipore), rabbit anti β--catenin (Ab2365. dilution: 1:5,000; Abcam), HA Tag antibody for SaCas9 protein (2-2.2.14, Thermofisher), rabbit anti-GFP (A11122, Invitrogen) followed by horseradish peroxidase-coupled antibodies (Goat anti-mouse: 115-035-146. dilution: 1:10,000 or Goat anti-Rabbit: 111-035-144. dilution: 1:50,000; Jackson ImmunoResearch). Blots were developed with ECL Plus reagents (Amersham Pharmacia) and imaged on the ChemiDoc Imaging System (BioRad).

### RNA-Seq Methods

Data from 4 LMI070 treated and 4 DMSO treated HEK293 cell groups were obtained after sequencing across two lanes run on an Illumina Hi-Seq 4000. The resulting fastq files were aligned to the GRCh38 human genome obtained from Ensembl using the STAR aligner.^19^ Splice junction output by STAR were quantified using a custom R script designed to identify splicing events unique to LMI070 treatment. Top ranking LMI070-exclusive splice events were manually assessed for their applicability to function as a splicing switch. A primary requirement of this evaluation was that two splice events (donor and acceptor) were identified as exclusive or enriched in LMI070 treated cells creating inclusion of a pseudoexon of reasonable size. Candidate splice events were visually evaluated using the Sashimi plot function available in IGV^20,21^. To assess the exclusivity of LMI070 induced splice sites to LMI070 treatment we evaluated the frequency with which candidate splice junctions were previously identified in diverse human RNA-Seq datasets deposited in the sequence read archive (SRA). This analysis was performed using Intropolis, a database of exon-exon junctions from 21,504 human RNA-Seq samples in the SRA archive. The Intropolis database is indexed by GRCh37 genomic position so we first converted our GRCh38 positions to GRCh37 using the LiftOver tool from the UCSC genome browser.^22^ We then queried LMI070 induced splice sites against the Intropolis database using a custom python script. The results for each LMI070 candidate splice event are summarized in Supplementary table 1.

Differential gene expression analysis was performed using DESeq2 to compare samples from the LMI070 and DMSO conditions.^23^ To visualize the abundance of meaningfully differentially expressed genes we generated a volcano plot with .05 Benjamini-Hochberg adjusted p-value threshold and a 0.1 log fold change threshold.

### Data availability

Custom R and Python scripts will be made available on Github.

RNA-Seq datasets will be archived in the NCBI Gene Expression Omnibus (GEO).

### Statistical analysis

Statistical analyses were performed using GraphPad Prism v5.0 software. Outlier samples were detected using the Grubb’s test (a = 0.05). Normal distribution of the samples was determined by using the D’Agostino and Pearson normality test. Data was analyzed using one-way ANOVA followed by a Bonferroni’s post hoc. Statistical significance was considered with a p < 0.05. All results are shown as the mean ± SEM.

## Acknowledgements

The authors would like to thank members of the Davidson lab and Ricardo Dolmetsch and Rajeev Sivasankaran of Novartis Institute for Biomedical Research for critical feedback and discussions, and the Hereditary Disease Foundation and the Children’s Hospital of Philadelphia Research Institute for support.

## Contributions

A.M.M. designed the research, performed experiments, analyzed data, and wrote the manuscript. A.A.H. and E.L. performed experiments. P.T.R. performed all bioinformatic analyses and wrote the manuscript. L.T. performed histology. A.M contributed to all *in vivo* studies. B.L.D designed and supervised the research, analyzed data, and wrote the manuscript.

