## Supplemental Material for "Regulated control of gene therapies with a drug induced switch"

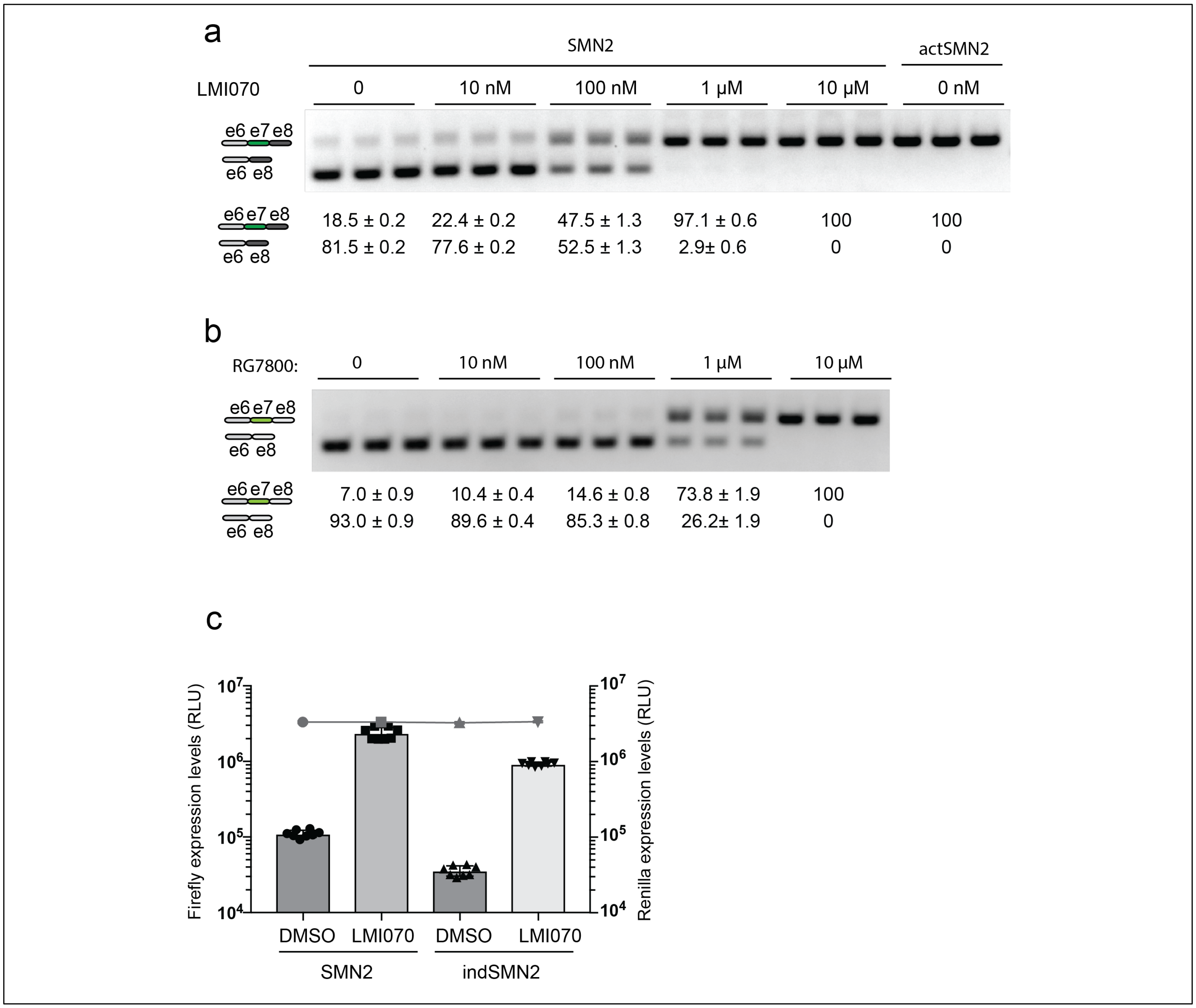


**ED Fig. 1a**. Exon 7 splicing of the SMN2 cassette in response to LMI070 dose. Representative RT-PCR reaction of exon 7 inclusion as a function of LMI070 dose. The quantification of the e7 spliced-in or spliced-out transcripts are the relative transcript levels presented as the mean ± SEM of 9 biological replicates.

**ED Fig. 1b**. Exon 7 splicing of the indSMN2 cassette in response to RG7800 dose. Representative RT-PCR reaction showing exon 7 inclusion as a function of RG7800 dose. The quantification of the e7 spliced-in or spliced-out transcripts are the relative transcript levels presented as the mean ± SEM of 8 biological replicates.

**ED Fig. F1c**. Luciferase activity of the SMN2 and indSMN2 cassettes in response to LMI070. Graph shows relative expression of Firefly luciferase expressed from the SMN2-on or indSMN2-on cassettes in cells treated with DMSO or LMI070 (100 nM). The activity of the transfection control *Renilla* luciferase cassette is represented as a line above the bar graph. The data are the mean ± SEM of 9 biological replicates.

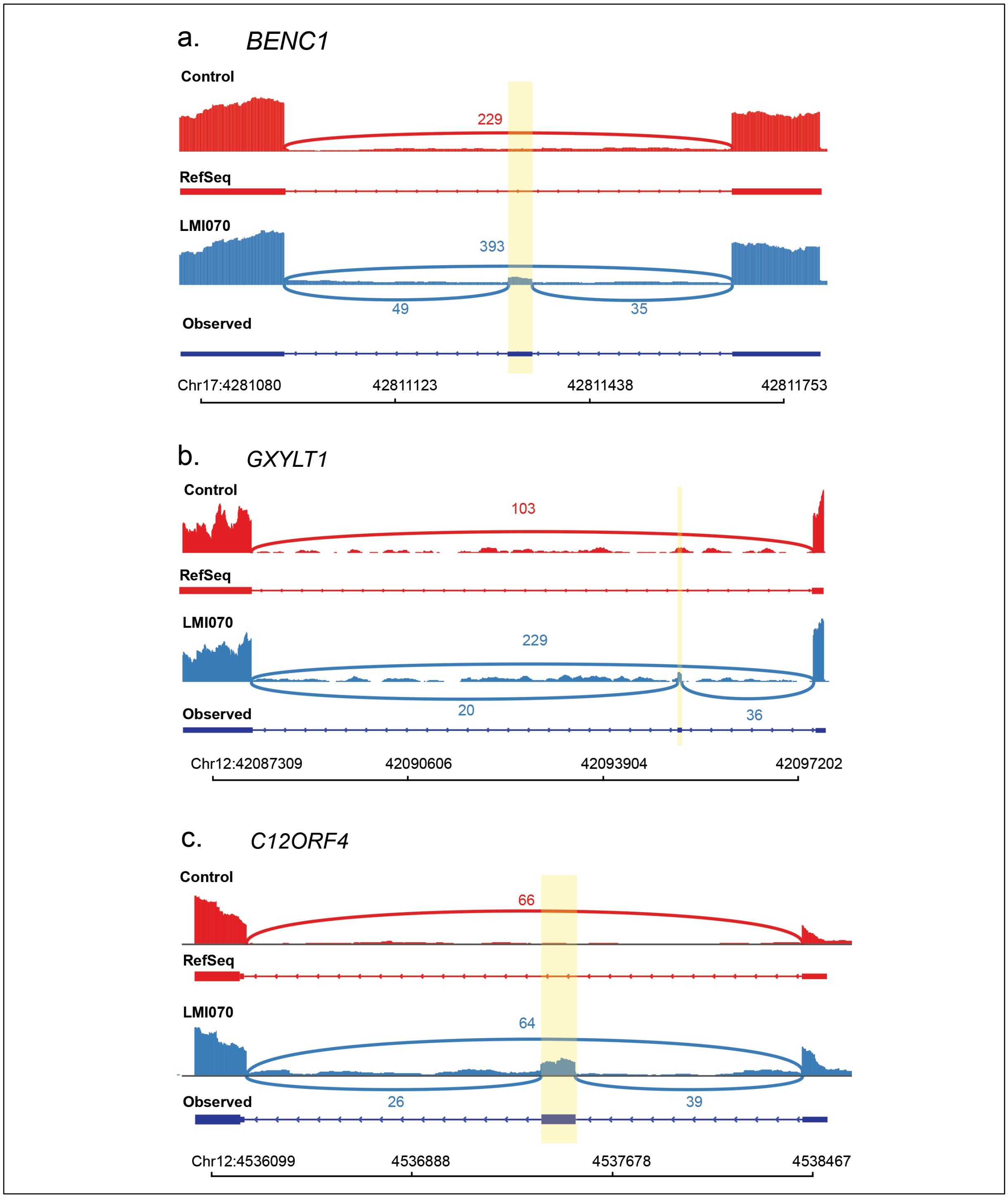


**ED Fig. 2a-c.** Sashimi plots depicting novel LMI070-spliced in exons for the top ranked genes identified by RNA-Seq. Genomic location and position of the LMI070 spliced in exon are indicated. (a) *BENC1*, (b) *GXYLT1*, (c) *C12ORF4*.

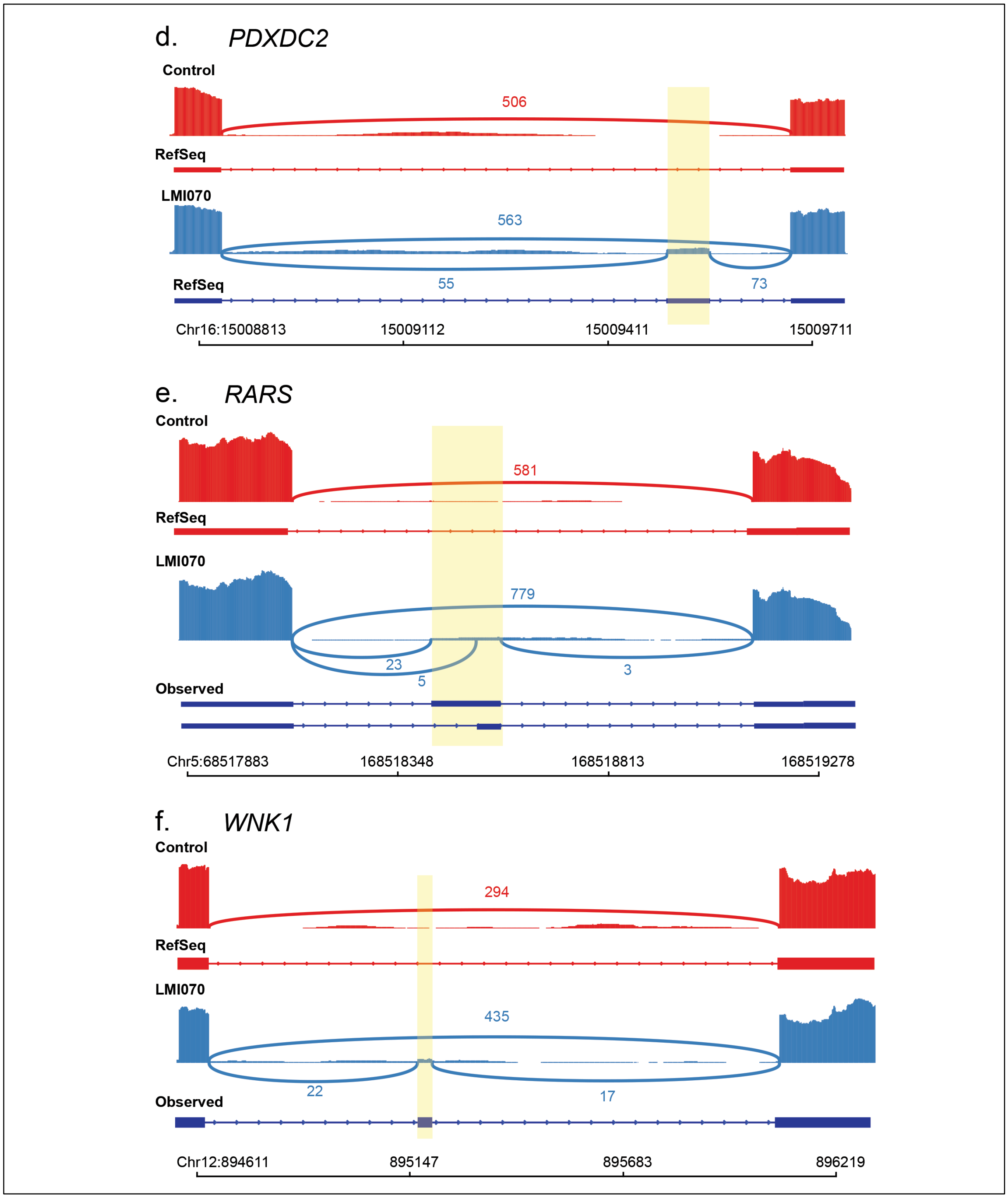


**ED Fig. 2d-f.** Sashimi plots depicting novel LMI070-spliced in exons for the top ranked genes identified by RNA-Seq. Genomic location and position of the LMI070 spliced in exon are indicated. (d) *PDXDC2*, (e) *RARS*, (f) *WNK1*.

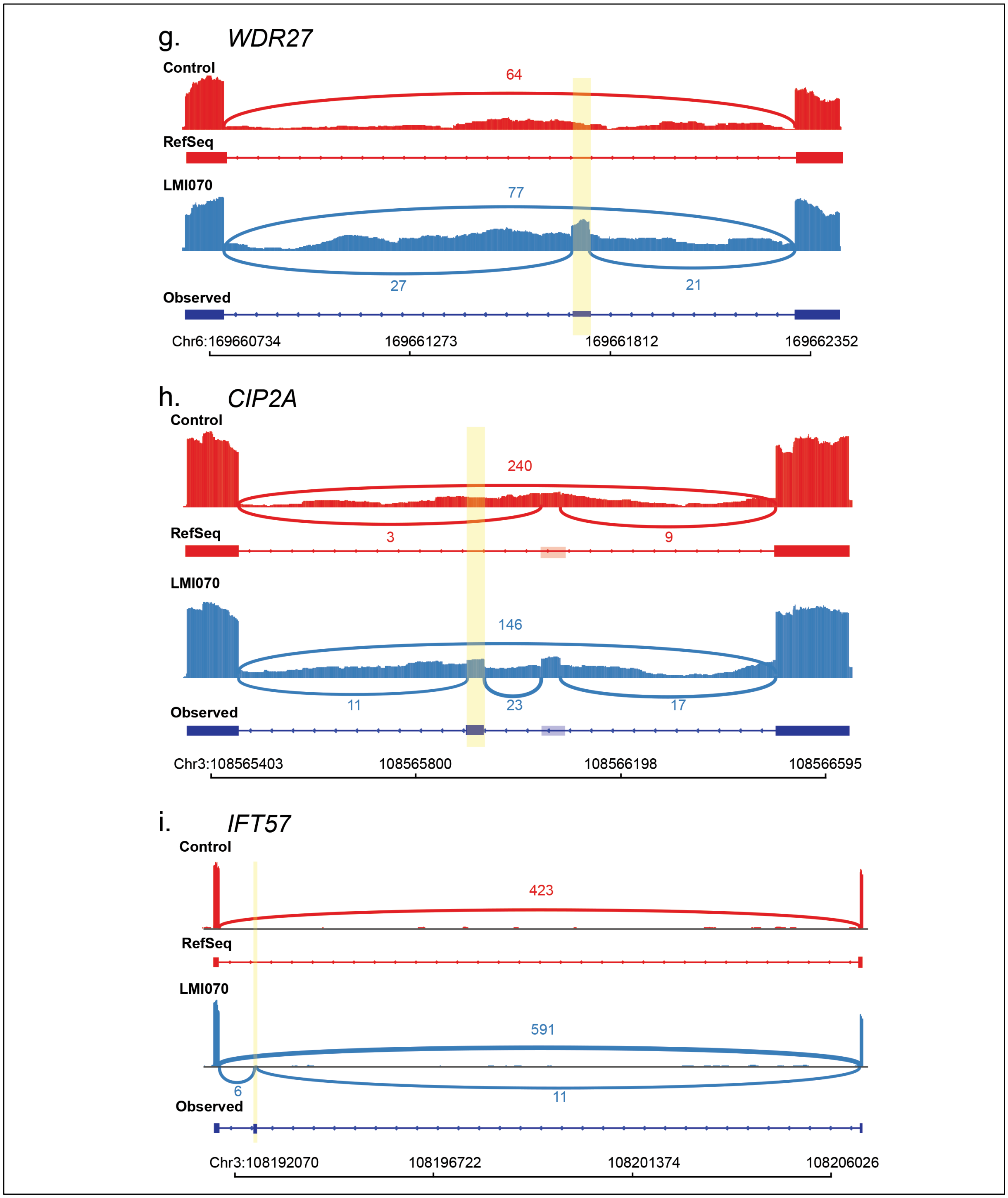


**ED Fig. 2g-i.** Sashimi plots depicting novel LMI070-spliced in exons for the top ranked genes identified by RNA-Seq. Genomic location and position of the LMI070 spliced in exon are indicated. (g) *WDR27*, (h) *CIP2A*, and (i) *IFT57*.

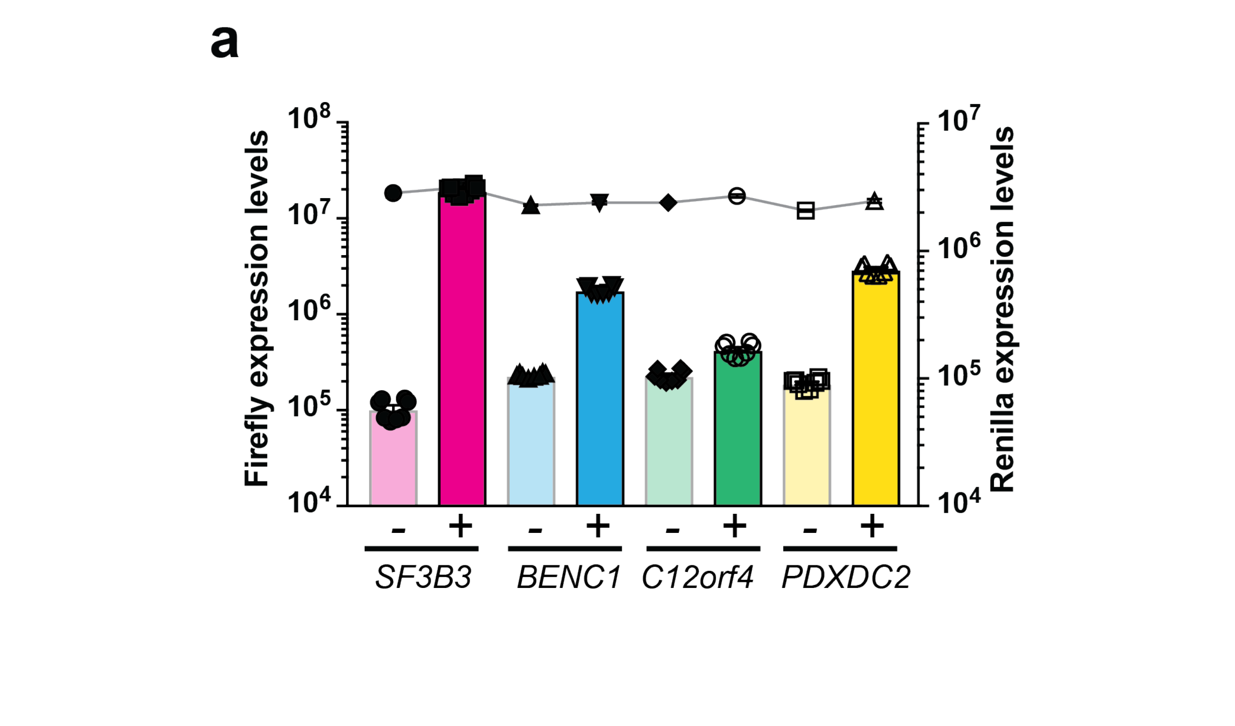


**ED Fig. 3a**: Luciferase activity of the minigene cassettes for *SF3B3, BENC1, C12ORF4 and PDXDC2*. Data show expression of *Firefly* luciferase from the minigenes in response to DMSO (depicted as minus) or LMI070 (depicted as plus) treatment relative to *Renilla* luciferase activity. Data are mean ± SEM of 8 biological replicates.

**b.** **Translation frequency from AUG and non-AUG codons determined from ribosome profiling.**

| Codons: | AUG>> | CUG>> | GUG> | ACG> | AUU= | AUA> | UUG= | AUC> | AAG= | AGG |
|---|---|---|---|---|---|---|---|---|---|---|
| Frequency: | 100 | 19 | 9 | 7 | 3 | 3 | 2 | 2 | 0.5 | 0.5 |

**Non-AUG codons in candidate minigene switches.**

*SF3B3*:

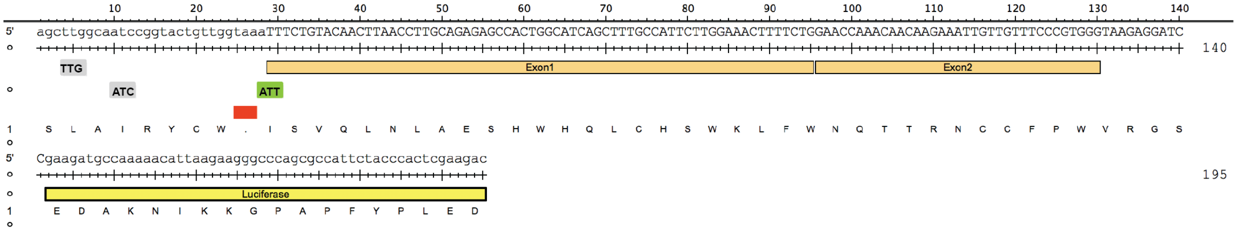


*BENC1:*

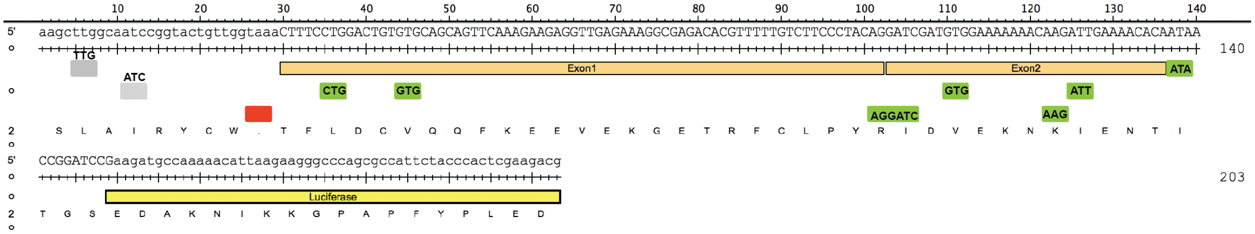


*PDXDC2*:

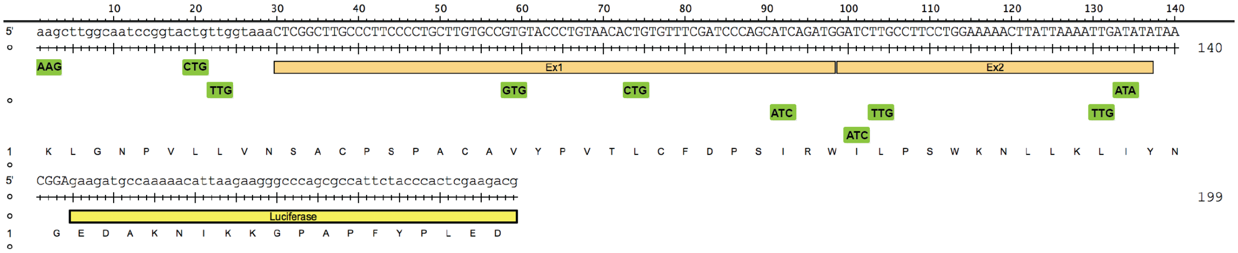


*C12ORF4*:

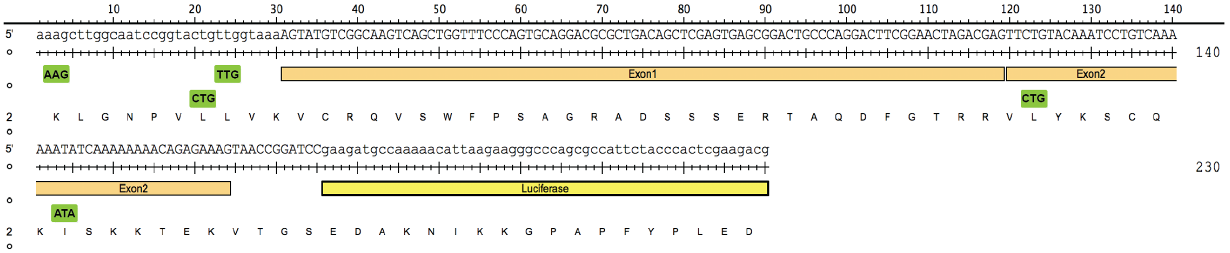


**ED Fig. 3b**. Depiction of the use frequency of the non-AUG start codons (CITE), and those in frame with the luciferase cDNA sequences in transcripts derived from the *SF3B3, BENC1, C12orf4 or PDXDC2* minigenes.

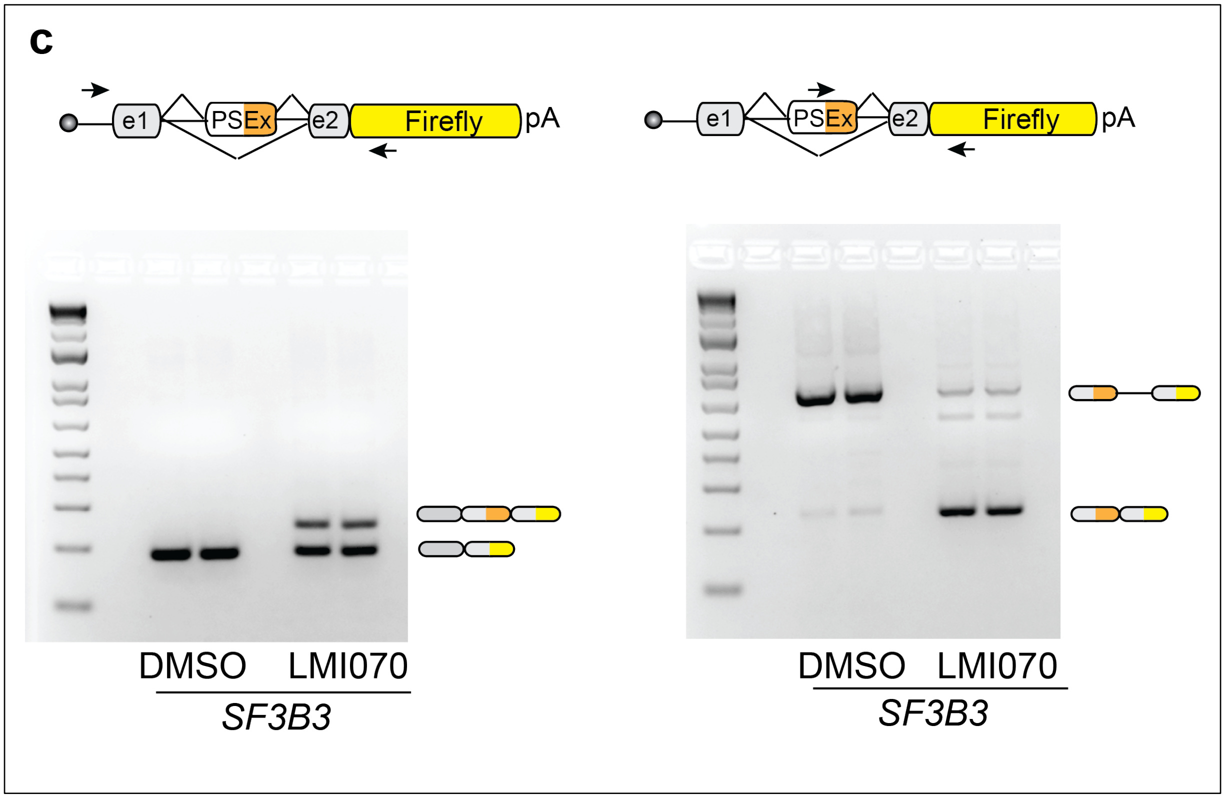


**ED Fig. 3c.** Representative RT-PCR reaction showing inclusion of the LMI070-induced *SF3B3* exon in response to DMSO or LMI070 treatment. Inclusion of the LMI070-spliced in exon was detected using primers binding the exons flanking the LMI070-induced exon (left), or using primers binding within the novel exon sequence (right).

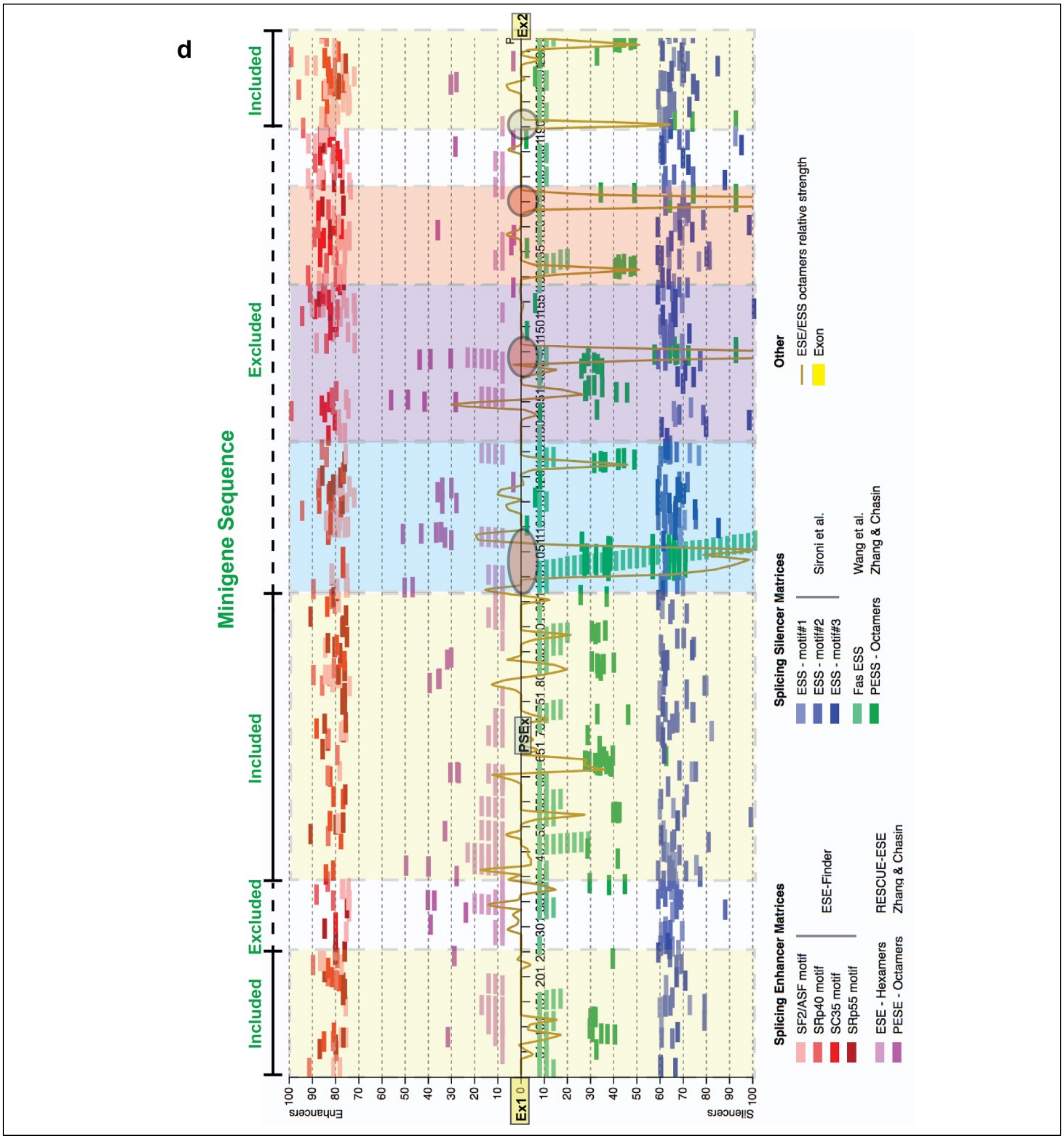


**ED Fig. 3d.** Graph depicting the position of predicted enhancer and silencer intronic sequences within the *SF3B3* intron. The elements were identified by the Human splicing finder website: (http://www.umd.be/HSF3/index.html). The position of the LMI070-induced exon (PSEx) and the intronic regions with a high density of silencer sequences are indicated. The grey and red shaded circles indicate intronic regions with a high density of silencer sequences contained with the *SF3B3* $X^{on}$ minigenes and the SF3B3int, SF3B3i1, SF3B3i2 and SF3B3i3 cassettes.

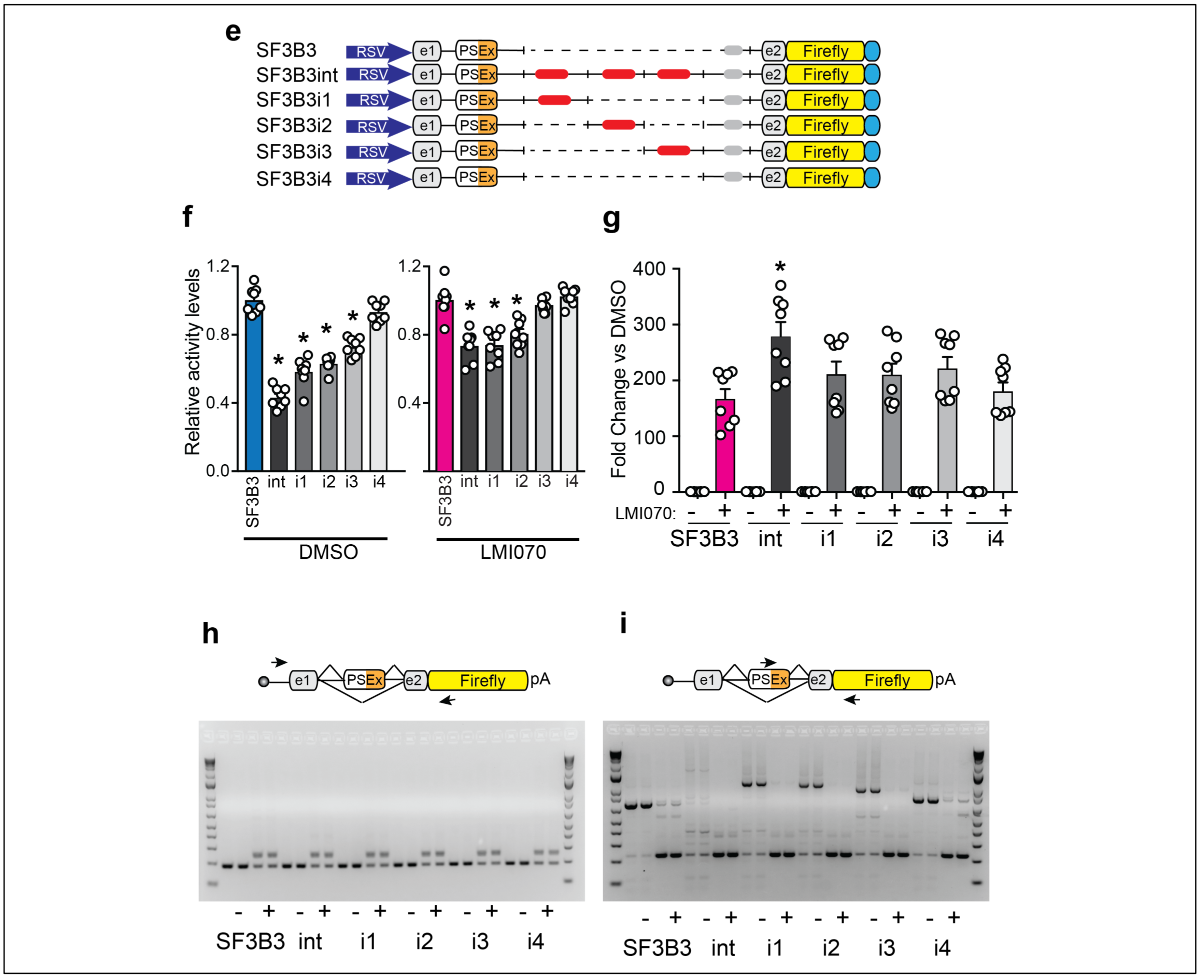


**ED Fig. 3e**. Cartoon depicting the *SF3B3* minigene cassettes containing the original minigene cassette (SF3B3), the full intron sequence (SF3B3int), versions with a high density of intronic silencer sequences (red circles; SF3B3i1, SF3B3i2, SF3B3i3), or a control sequence lacking these same silencers sequences (SF3B3i4).

**ED Fig. 3f.** Luciferase activity of various *SF3B3* minigene constructs containing additional *SF3B3* intron regions. HEK293 cells were transfected with equimolar amounts of plasmids and luciferase activity was determined 24h after transfection. The graphs show luciferase activity of new SF3B3 cassettes after DMSO (left) or LMI070 (right) treatment and are relative to the original SF3B3 minigene switch (blue and pink for DMSO or LMI070, respectively). Data are the mean ± SEM of 8 biological replicates (*p < 0.0001, one-way ANOVA followed by a Bonferroni's post hoc).

**ED. Fig. 3g.** Fold-induction of luciferase of the *SF3B3* minigene constructs. The fold change of luciferase activity in LMI070-treated samples is relative to DMSO-treated cells. All samples are normalized to *Renilla* luciferase activity. The original SF3B3 minigene switch is denoted (pink). Data are the mean ± SEM of 8 biological replicates (*p < 0.0001, one-way ANOVA followed by a Bonferroni's post hoc).

**ED Fig. 3h,i.** Representative gels showing PCR assay for the LMI070-induced pseudoexon. Priming was either to the exons flanking the pseudoexon (h), or within the pseudoexon sequence (i).

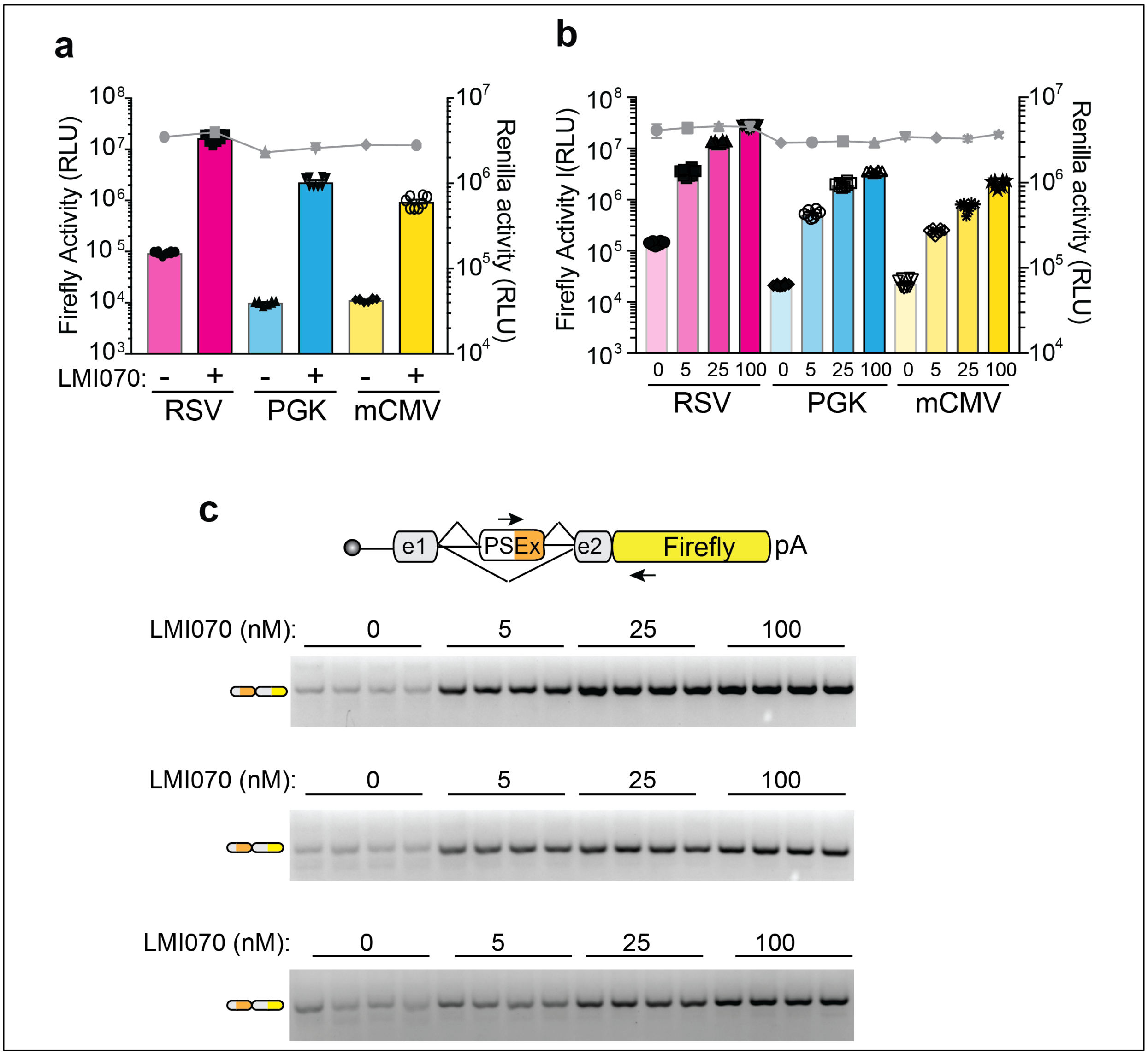


**ED Fig. 4a, b.** Activity of the SF3B3-X$^{on}$ cassette when expressed various promoters and their responsiveness to LMI070 dose. (a) *Firefly* luciferase from the X$^{on}$ cassettes in response to DMSO or LMI070 treatment (minus, plus, respectively) relative to *Renilla* luciferase (grey line). (b) *Firefly* luciferase from the X$^{on}$ cassettes in response to varying doses of LMI070 relative to *Renilla* luciferase (grey line). The data are the mean ± SEM of 8 biological replicates.

**ED Fig. 4c.** Representative gels from RT-PCR analysis for assessment of the LMI070-induced pseudoexons expressed from the noted promoters in response varying doses of LMI070. Pseudoexon inclusion was detected using primers binding within the LMI070-induced pseudoexon and the downstream exon. Splicing was quantified and transcript levels presented as the mean ± SEM of 8 biological replicates.

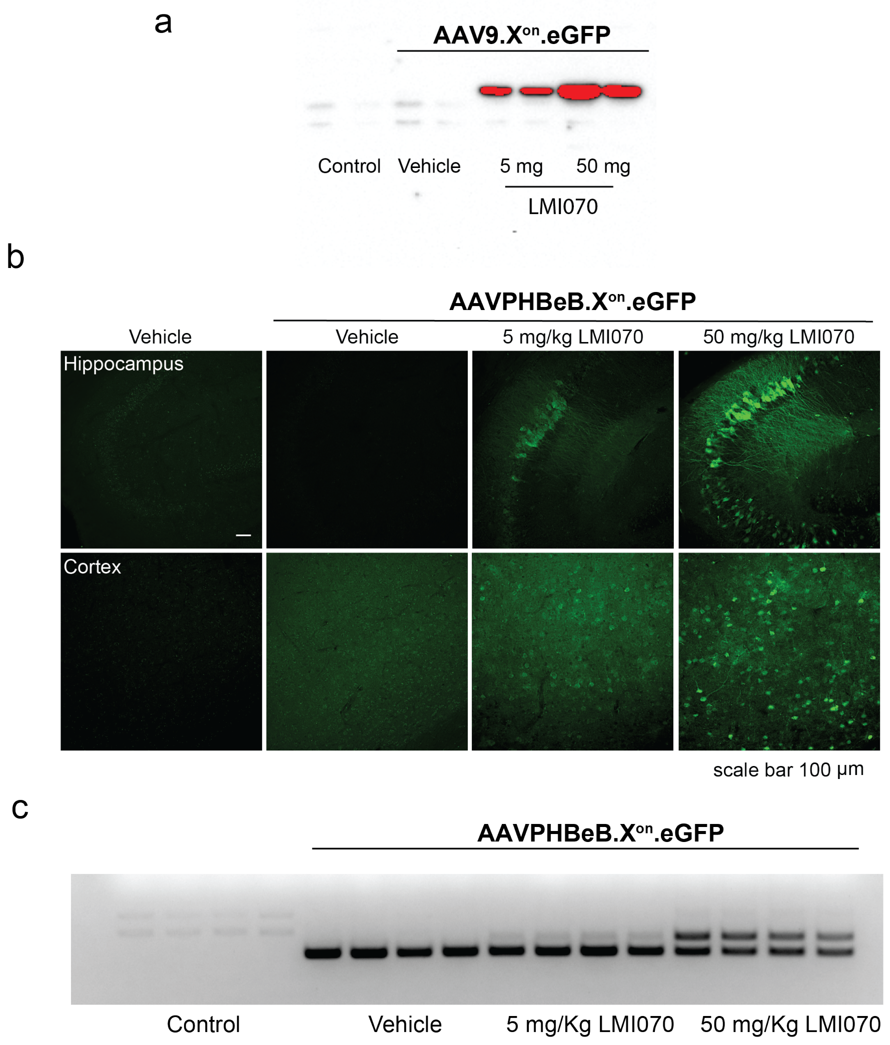


| | Control | Vehicle | 5 mg/Kg LMI070 | 50 mg/Kg LMI070 |
|---|---|---|---|---|
| AVG Ct eGFP | ND | 25.96 ± 0.58 | 26.67 ± 0.48 | 27.02 ± 0.81 |
| AVG Ct XS | ND | 38.29 ± 0.77 | 31.39 ± 0.36 | 28.65 ± 0.87 |
| Fold induction | ND | 1 | 182 | 1508 |

**ED Fig. 5a.** Extended exposure of the western blot from Fig. 5c.
**ED Fig. 5b.** Photomicrographs of tissue sections showing eGFP expression in brain from mice treated iv 4 weeks earlier with AAVPHBeB.$X^{on}$.eGFP, and 24h after treatment with LMI070 at 5 or 50 mg/kg. eGFP in hippocampus and cortex are shown (scale bar 100 μm).
**ED Fig. 5c.** Agarose gel from PCR assays demonstrates inclusion of the splicing activity after AAVPHBeB.$X^{on}$.eGFP gene delivery, in response LMI070 (n=4). Data shows the average Ct values for eGFP or LMI070-induced expression using the $X^{on}$ gene expression assays depicted in d). Fold change of the spliced expression cassette is shown relative to basal levels in mice injected with AAVPHBeB.$X^{on}$.eGFP and treated with vehicle.

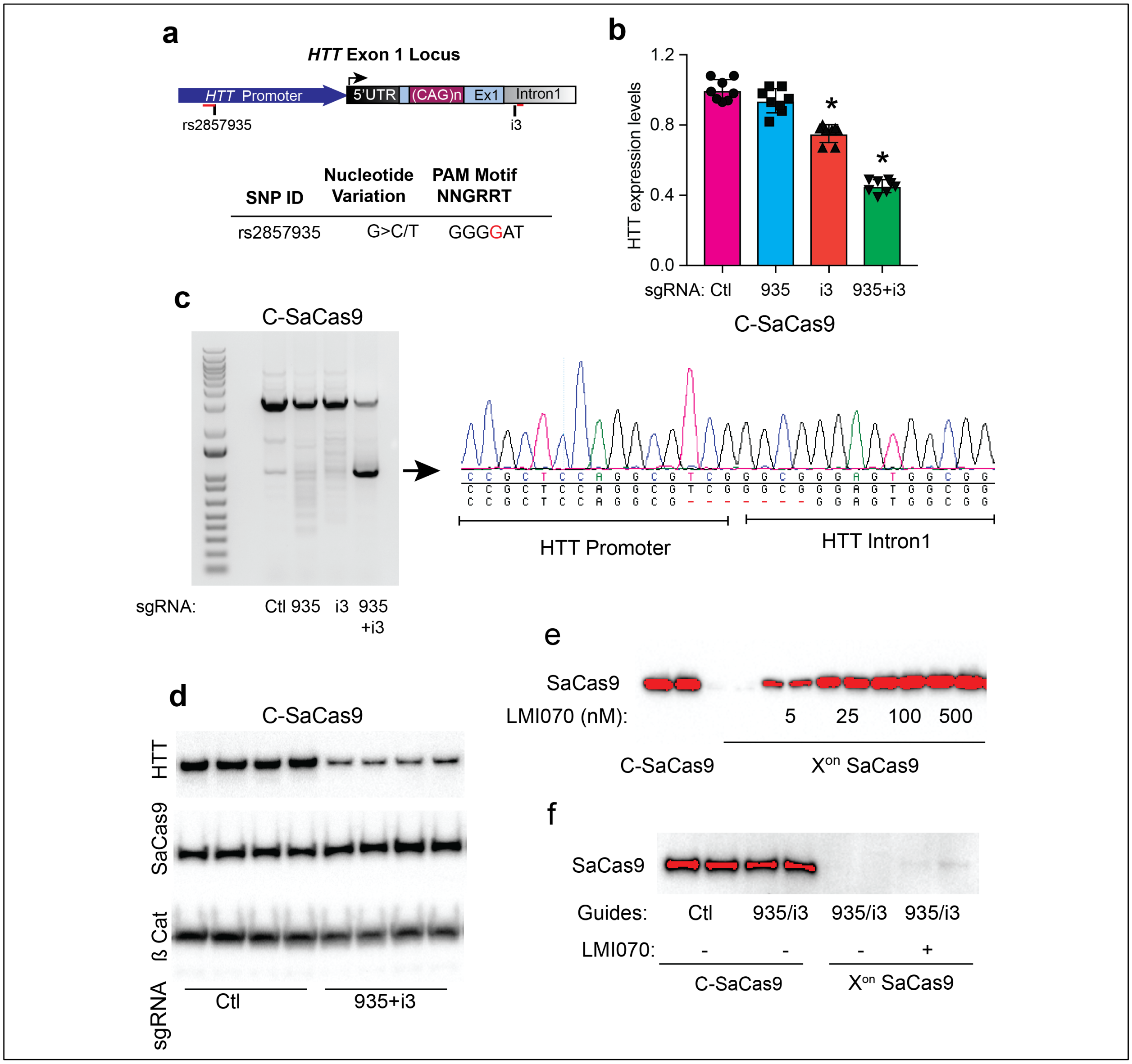


**ED Fig. 6a**. Cartoon depicting the allele-specific editing strategy for targeted deletions of mutant *HTT* exon 1. SNPs within SaCas9 PAM sequences upstream of HTT exon-1 allow selective editing of the mutant allele when present in heterozygosity. After DNA repair, mutant HTT exon-1 is removed by a pair of sgRNA/SaCas9 complexes binding upstream and downstream of exon-1. The SNP (referred to as 935) is annotated.
**ED Fig. 6b**. HTT mRNA levels in HEK293 cells transfected with constitutive SaCas9 CRISPR cassettes targeting *HTT* exon 1 flanking sequences as assessed by RT-qPCR. Samples are normalized to human GAPDH, and data are the mean ± SEM relative to cells transfected with SaCas9 CRISPR expression plasmids together with the control sgRNA sequence (n = 8 biological replicates; *p < 0.0001, one-way ANOVA followed by Bonferroni's post hoc).

**ED Fig. 6c.** Representative gel showing HTT exon-1-targeted deletion in cells transfected with SaCas9 CRISPR expression plasmids together with the control sgRNA sequence or sgRNA sequences targeting the flanking sequences of HTT exon 1 and the selective reporter eGFP/puromycin expression cassettes. The Sanger sequencing results of the small PCR products confirmed deletion of HTT exon 1 at the DNA cleavage site.
**ED Fig. 6d.** A representative western blot showing HTT exon1 levels in cells transfected with SaCas9 CRISPR plasmids and the noted sgRNAs. SaCas9 and β catenin was used as protein loading controls, respectively.
**ED Fig. 6e.** Extended exposure of SaCas9 expression of Fig. 6b.
**ED Fig. 6f**. Extended exposure of SaCas9 expression from Fig. 6g.

**Supplementary Table 1.**
The genomic locations of candidate LMI070-induced exons and the frequency of events from RNA-Seq datasets. Summarized here are the differentially expressed candidate LMI070-induced splicing positions as identified by RNA-Seq of HEK293 cells treated with either DMSO or LMI070 (25nM). All candidates shown were manually selected from a bioinformatically generated list of top hits based on their suitability for construction of an exon switching genomic minigene, their exclusivity to the LMI070 condition, and their minimal to undetectable levels in DMSO treated cells. The top 25 rows (shaded green) indicate hits observed exclusively upon LMI070 exposure. The following 22 rows (shaded yellow) indicate candidates where splicing was enriched but not totally exclusive to LMI070 treatment. Columns (from left to right) indicate: 1. Exclusivity to LMI070 induction. 2. The gene ID containing the splicing events of interest. 3. The GRCh38 genomic positions used to create the splice event-containing genomic minigene. 4. The GRCh38 genomic positions of the pseudoexon created by LMI070-induced splicing. 5. The number of exon-exon junction spanning reads observed with DMSO treatment. 6. The number of exon-exon junctions spanning reads observed with LMI070 treatment. To assess the frequency with which LMI070-induced events occur we queried Intropolis[1], a database containing the frequency of splicing events observed across 21,504 human RNA-Seq samples, representing a diverse set of human tissues and conditions. The reference genome used for the Intropolis database is GRCh37 so the LiftOver feature from the UCSC genome browser was used to convert the GRCh38 coordinates to GRCh37. Column 7. The GRCh37 genomic position of the minigene. 8. The GRCh37 genomic position of the pseudoexon. 9. The DNA sequence of the LMI070 binding sequence in the pseudoexon. 10. The position of canonical splice junction (CJ). 11. The number of Intropolis RNA-Seq datasets in which each canonical splice event was observed. 12. The total number of observations identified for each canonical splice site. 13. The position of junction 1 (J1) and the first LMI070-induced exon-exon junction (sorted by genomic position) connecting a canonical exon to a LMI070-induced pseudoexon. 14 and 15 indicate the number and percentage of Intropolis datasets in which each S1 splice event was observed. Columns 16 and 17 indicate the number and percentage of total counts in which each S1 splice event was observed. 18. The position of junction 2 (J2) the second LMI070 induced exon-exon junction (sorted by genomic position) connecting a LMI070 induced pseudoexon to a canonical exon. Columns 19 and 20 list the number and percentage of

Intropolis datasets in which the LMI070 induced splicing event was observed. Columns 21 and 22 indicate the total number and percentage of reads containing each junction in the Intropolis dataset.

1 Nellore, A. *et al.* Human splicing diversity and the extent of unannotated splice junctions across human RNA-seq samples on the Sequence Read Archive. *Genome Biol* **17**, 266, doi:10.1186/s13059-016-1118-6 (2016).

### ED Table 1

| | | GRCh38 | | | |
|---|---|---|---|---|---|
| | **Gene ID** | **Genomic Minigene region** | **Pseudo exon position** | **DMSO Observed Avg Counts (J1, J2)** | **LMI070 Observed Avg Counts (J1, J2)** |
| **Exclusive** | *SF3B3* | chr16:70,526,657-70,529,199 | chr16:70,527,376-70,527,429 | 0, 0 | 31, 30.5 |
| | *BENC1* | chr17:42,810,759-42,811,797 | chr17:42,811,292-42,811,330 | 0, 0 | 24.45, 15.75 |
| | *GXYLT1* | chr12:42,087,786-42,097,614 | chr12:42,095,151-42,095,214 | 0, 0 | 10.75, 23.75 |
| | *SKP1* | chr5:134,173,809-134,177,053 | chr5:134,175,284-134,175,385 | 0, 0 | 5.75, 23.75 |
| | *SKP1* | chr5:134,173,809-134,177,053 | chr5:134,175,284-134,175,423 | 0, 0 | 15.25, 11.75 |
| | *C12orf4* | chr12:4,536,017-4,538,508 | chr12:4,537,380-4,537,514 | 0, 0 | 17.5, 19 |
| | *SSBP1* | chr7:141,739,167-141,742,229 | chr7: 141,741,310-141,741,459 | 0, 0 | 17.25, 2.25 |
| | *RARS* | chr5:168,517,815-168,519,190 | chr5:168,518,369-168,518,523 | 0, 0 | 13.5, 1.5 |
| | *RARS* | chr5:168,517,815-168,519,190 | chr5:168,518,469-168,518,523 | 0, 0 | 13.6, 1.75 |
| | *PDXDC2P* | chr16:70,030,988-70,031,968 | chr16:70,031,186-70,031,248 | 0, 0 | 13.25, 10.25 |
| | *STRADB* | chr2:201,469,953-201,473,076 | chr2:201,470,907-201,471,111 | 0, 0 | 9.5, 5.25 |
| | *WNK1* | chr12:894,562-896,732 | chr12:895,161-895,196 | 0, 0 | 9, 5.5 |
| | *WDR27* | chr6:169,660,663-169,662,424 | chr6:169,661,703-169,661,750 | 0, 0 | 8.5, 7.25 |
| | *CIP2A* | chr3:108,565,355-108,566,638 | chr3:108,565,898-108,565,931 | 0, 0 | 7.75, 5 |
| | *ITF57* | chr3:108,191,521-108,206,696 | chr3:108,192,476-108,192,526 | 0, 0 | 7.25, 5.25 |
| | *HTT* | chr4:3,212,555-3,214,145 | chr4:3213622-3213736 | 0, 0 | 7, 2.25 |
| | *SKA2* | chr17:59,112,228-59,119,514 | chr17:59119395-59119495 | 0, 0 | 6.75, 1 |
| | *EVC* | chr4:5,733,318-5,741,822 | chr4:5741334-5741441 | 0, 0 | 6.5, 3 |
| | *DYRK1A* | chr21:37,420,144-37,473,056 | chr21:37422581-37422652 | 0, 0 | 6.25, 6 |
| | *GNAQ* | chr9:77,814,652-77,923,557 | chr9:77920648-77920703 | 0, 0 | 6, 1 |
| | *ZMYM6* | chr1:35,019,257-35,020,472 | chr1:35020261-35020279 | 0, 0 | 5.75, 4 |
| | *CYB5B* | chr16:69,448,031-69,459,260 | chr16:69448605-69448753 | 0, 0 | 5.75, 1.25 |
| | *MMS22L* | chr6:97,186,342-97,229,533 | chr6:97201362-97201465 | 0, 0 | 5.75, 2 |
| | *MEMO1* | chr2:31,883,262-31,892,301 | chr2:31,887,035-31887087 | 0, 0 | 5, 2.25 |
| | *PNISR* | chr6:99,416,278-99,425,413 | chr6:99420523-99420584 | 0, 0 | 5, 4 |
| **Enriched** | *CACNA2D1* | chr7:82,066,406-82,084,958 | chr7:82,076,016-82,076,122 | 0.25, 0.75 | 18.5, 1.5 |
| | *SSBP1* | chr7:141,739,083-141,742,248 | chr7:141741310-141741459 | 0.25, 0 | 16.75, 2.25 |
| | *DDX42* | chr17:63,805,048-63,806,672 | chr17:63,806,151-63,805,994 | 0.25, 0 | 13.75, 4.5 |
| | *ASAP1* | chr8:130,159,817-130,167,688 | chr8:130,160,785-130,160,793 | 0.25, 0.25 | 13, 11.5 |
| | *DUXAP10* | chr14:19,294,564-19,307,199 | chr14:19,305,354-19,305,469 | 0.25, 0.75 | 9.5, 1.25 |
| | *AVL9* | chr7:32,558,783-32,570,372 | chr7:32,562,558-32,562,913 | 0.25, 0.5 | 7.5, 1.5 |
| | *DYRK1A* | chr21:37,419,920-37,472,960 | chr21:37,422,582-37,422,652 | 0.25, 0 | 6.25, 6 |
| | *FAM3A* | chrX:154,512,311-154,512,939 | chrX: 154,512,568-154,512,706 | 0.25, 0 | 5.75, 1 |
| | *FHOD3* | chr18:36,740,620-36,742,886 | chr18:36,742,377-36,742,468 | 0.5, 0.25 | 15.25, 3.5 |
| | *TBCA** | chr5:77,707,994-77,777,000 | chr5:77,774,217 | 0.5 | 14 |
| | *MZT1* | chr13:72,718,939-72,727,611 | chr13:72,725,642-72,725,778 | 0.5, 1 | 13.25, 1.25 |
| | *LINC01296* | chr14:19,092,877-19,096,652 | chr14:19094556-19094671 | 0.5, | 8.25, |
| | *SF3B3* | chr16:70,541,627-70,544,553 | chr16:70544169-70544249 | 0.5, 0 | 8.25, 3 |
| | *SAFB* | chr19:5,654,060-5,654,457 | chr19:5,654,140-5,654,368 | 0.5, 0 | 6.25, 2 |
| | *GCFC2* | chr2:75,702,163-75,706,652 | chr2:75,702,691-75,702,807 | 0.5, 0 | 6.25, 2 |
| | *MRPL45* | chr17:38,306,450-38,319,088 | chr17:38,312,587-38,312,661 | 0.5, 0 | 5.75, 1.25 |
| | *SPIDR* | chr8:47,260,788-47,280,196 | chr8:47,273,337-47,273,450 | 0.5, 0 | 5.5, 1.75 |
| | *DUXAP8* | chr22:15,815,315-15,828,713 | chr22:15,817,119-15,817,234 | 0.75, 0.25 | 13.75, 1.25 |
| | *PDXDC1* | chr16:15,008,772-15,009,763 | chr16:15,009,499-15,009,561 | 2, 1 | 16.75, 1.5 |
| | *MAN1A2* | chr1:117,442,104-117,461,030 | chr1:117,456,085-117,456,206 | 0.75, 1 | 8, 5.25 |
| | *RAF1* | chr3:12,600,376-12,604,350 | chr3:12,603,478-12,603,537 | 1, 0.25 | 16.5, 1 |
| | *ERGIC3* | chr20:35,548,787-35,554,452 | chr20:35,549,163-35,549,207 | 1, 0 | 7.5, 2.5 |

| | | | Canonical Junction Metrics | | | GRCh37 (hg1 |
|---|---|---|---|---|---|---|
| Genomic Minigene region | Pseudo exon position | MI070 Binding sequen | CJ exon-exon juction | CJ Intropolis Datasets | CJ_TotalCounts | J1 Position |
| chr16:70560560-70563102 | chr16:70561279-70561332 | AGAGTAAGAC | chr16:70560630-70562775 | 12873 | 825337 | chr16:70560630-70561278 |
| chr17:40962777-40963815 | chr17:40963310-40963348 | AGAGTAAGGC | chr17:40962947-40963672 | 13635 | 910210 | chr17:40962947-40963309 |
| chr12:42481588-42491416 | chr12:42488953-42489016 | AGAGTATAGT | chr12:42481750-42491243 | 7430 | 43383 | chr12:42481750-42488952 |
| chr5:133509500-133512744 | chr5:133510975-133511076 | AGAGTAGGAT | chr5:133509714-133512545 | 13277 | 2756168 | chr5:133509714-133510974 |
| chr5:133509500-133512744 | chr5:133510975-133511114 | AGAGTAGGAT | chr5:133509714-133512545 | 13277 | 2756168 | chr5:133509714-133510974 |
| chr12:4645183-4647674 | chr12:4646546-4646680 | AGAGTAAGAA | chr12:4645386-4647574 | 9856 | 102163 | chr12:4645386-4646545 |
| chr7:141438967-141442029 | chr7:141441110-141441259 | AGAGTAAGGC | chr7:141438991-141441968 | 14501 | 2473164 | chr7:141438991-141441109 |
| chr5:167944820-167946195 | chr5:167945374-167945528 | AGAGTAGGAT | chr5:167945068-167946085 | 13602 | 826669 | chr5:167945068-167945373 |
| chr5:167944820-167946195 | chr5:167945474-167945528 | AGAGTAGGAT | chr5:167945068-167946085 | 13602 | 826669 | chr5:167945068-167945473 |
| chr16:70064891-70065871 | chr16:70065089-70065151 | AGAGTAAGAA | chr16:70064970-70065802 | 9586 | 117767 | chr16:70064970-70065088 |
| chr2:202334676-202337799 | chr2:202335630-202335834 | AGAGTAAGGA | chr2:202334776-202337677 | 9420 | 155599 | chr2:202334776-202335629 |
| chr12:1003728-1005898 | chr12:1004327-1004362 | AGAGTAGGTG | chr12:1003802-1005236 | 12935 | 800572 | chr12:1003802-1004326 |
| chr6:170060759-170062520 | chr6:170061799-170061846 | AGAGTAAGCA | chr6:170060863-170062399 | 9937 | 128203 | chr6:170060863-170061798 |
| chr3:108284202-108285485 | chr3:108284745-108284778 | AGAGTAAGAA | chr3:108284302-108285343 | 9210 | 124822 | chr3:108284302-108284744 |
| chr3:107910368-107925543 | chr3:107911323-107911373 | AGAGTAGGCC | chr3:107910491-107925474 | 12662 | 583056 | chr3:107910491-107911322 |
| chr4:3214282-3215872 | chr4:3215349-3215463 | AGAGTAAGGG | chr4:3214437-3215684 | 11146 | 243427 | chr4:3214437-3215348 |
| chr17:57189589-57196875 | chr17:57196756-57196856 | AGAGTAAGAG | chr17:57189707-57196679 | 10377 | 297301 | chr17:57189707-57196756 |
| chr4:5735045-5743549 | chr4:5743061-5743168 | AGAGTAAGCA | chr4:5735163-5743442 | 6716 | 74348 | chr4:5735163-5743060 |
| chr21:38792446-38845358 | chr21:38794883-38794954 | AGAGTAGGTT | chr21:38792687-38844985 | 10821 | 152058 | chr21:38792687-38794883 |
| chr9:80429568-80538473 | chr9:80535564-80535619 | AGAGTAAGCT | chr9:80430687-80537076 | 10041 | 224120 | chr9:80430687-80535563 |
| chr1:35484858-35486073 | chr1:35485862-35485880 | ACTGTGAGTA | chr1:35485204-35485983 | 11692 | 199379 | chr1:35485204-35485861 |
| chr16:69481934-69493163 | chr16:69482508-69482656 | TAGGTGGTTC | chr16:69482048-69492995 | 13834 | 1150919 | chr16:69482048-69482509 |
| chr6:97634218-97677409 | chr6:97649238-97649341 | GAGGTGATTG | chr6:97634567-97676769 | 8959 | 91164 | chr6:97634567-97649237 |
| chr2:32108331-32117370 | chr2:32112104-32112156 | AGAGTAAGGT | chr2:32108532-32117060 | 8678 | 99228 | chr2:32108532-32112103 |
| chr6:99864154-99873289 | chr6:99868399-99868460 | AGAGTAGTGT | chr6:99864305-99873090 | 13184 | 888372 | chr6:99864305-99868398 |
| chr7:81695722-81714274 | chr7:81705332-81705438 | CAGGTTGGTA | chr7:81695841-81714084 | 6256 | 96528 | chr7:81695841-81705331 |
| chr7:141438883-141442048 | chr7:141441110-141441259 | AGAGTAAGGC | chr7:141438991-141441968 | 14501 | 2473164 | chr7:141438991-141441109 |
| chr17:61882408-61884032 | Unable to lift over | AGAGTAAGAT | chr17:61882536-61883894 | 12718 | 722896 | NA |
| chr8:131172063-131179934 | chr8:131173031-131173039 | AGAGTAAGTA | chr8:131172211-131179781 | 11448 | 396871 | chr8:131172211-131173030 |
| chr14:19882243-19894878 | chr14:19893035-19893150 | AGAGTAAGGT | chr14:19884030-19894699 | 638 | 1502 | Not Found |
| chr7:32598395-32609984 | chr7:32602170-32602525 | AGAGTAAGAC | chr7:32599077-32609631 | 11339 | 199392 | chr7:32599077-32602169 |
| chr21:38792222-38845262 | chr21:38794884-38794954 | AGAGTAGGTT | chr21:38792687-38844985 | 10821 | 152058 | chr21:38792687-38794883 |
| chrX:153740635-153741263 | chrX:153740892-153741030 | GGGGTAGGGA | chrX:153740736-153741146 | 9459 | 98652 | chrX:153740736-153740891 |
| chr18:34320583-34322849 | chr18:34322340-34322431 | AGAGTAAGAG | chr18:34320802-34322699 | 7279 | 113308 | chr18:34320802-34322339 |
| chr5:77003819-77072824 | chr5:77070041 | ND | chr5:77004173-77072028 | 14180 | 942768 | NA |
| chr13:73293077-73301749 | chr13:73299780-73299916 | AGAGTAAGAA | chr13:73293236-73301661 | 12409 | 516948 | chr13:73293236-73299779 |
| chr14:19680556-19684333 | chr14:19682237-19682352 | AGAGTAAGAT | chr14:19680686-19683691 | 239 | 315 | chr14:19680686-19682236 |
| chr16:70575530-70578456 | chr16:70578072-70578152 | AGAGTAAAGA | chr16:70575738-70578340 | 12928 | 880623 | chr16:70575738-70578071 |
| chr19:5654071-5654468 | chr19:5654151-5654379 | AGAGTAAGGA | chr19:5654212-5654378 | 13168 | 1083231 | Not Found |
| chr2:75929289-75933778 | chr2:75929817-75929933 | TGAGTAAGAG | chr2:75929550-75933648 | 11225 | 168121 | chr2:75929550-75929816 |
| chr17:36462417-36474972 | chr17:36468550-36468624 | AGAGTAAGAC | chr17:36462599-36474585 | 12718 | 608565 | chr17:36462599-36468549 |
| chr8:48173380-48192784 | chr8:48185929-48186042 | AGAGTAAGAC | chr8:48173584-48192449 | 10729 | 158491 | chr8:48173584-48185928 |
| chr14:19,680,685-19,691,354 | chr14:19682237-19682352 | AGAGTAAGAT | chr14:19680686-19683691 | 239 | 315 | chr14:19680686-19682236 |
| chr16:15102629-15103620 | chr16:15103356-15103418 | AGAGTAAGAA | chr16:15102705-15103537 | 12939 | 767189 | chr16:15102705-15103355 |
| chr1:117984726-118003652 | chr1:117998707-117998828 | AGAGTAAGGT | chr1:117984948-118003110 | 11582 | 229559 | chr1:117984948-117998706 |
| chr3:12641875-12645849 | chr3:12644977-12645036 | AGAGTAGGTA | chr3:12641915-12645634 | 12808 | 832362 | chr3:12641915-12644976 |
| chr20:34136540-34142223 | chr20:34136917-34136961 | GTGGTAGGTA | chr20:34136619-34142142 | 4423 | 84287 | chr20:34136619-34136916 |

.9)

| LMI070 Junction1 Metrics | | | | LMI070 Junction2 Metrics | | | | |
|---|---|---|---|---|---|---|---|---|
| J1 NumDatasets | J1 % Data Sets | J1 TotalCounts | J1 % Total counts | J2 Positon | J2 NumDatasets | J2 % Data Sets | J2 TotalCounts | J2 % Total counts |
| 10 | 0.08 | 13 | 0.002 | chr16:70561333-70562775 | 1 | 0.01 | 1 | 0.0001 |
| 301 | 2.21 | 552 | 0.061 | chr17:40963349-40963672 | 182 | 1.33 | 303 | 0.0333 |
| 8 | 0.11 | 9 | 0.021 | chr12:42489017-42491243 | 79 | 1.06 | 97 | 0.2236 |
| 15 | 0.11 | 16 | 0.001 | chr5:133511077-133512545 | 6 | 0.05 | 6 | 0.0002 |
| 15 | 0.11 | 16 | 0.001 | chr5:133511115-133512545 | 13 | 0.10 | 18 | 0.0007 |
| 354 | 3.59 | 563 | 0.551 | chr12:4646681-4647574 | 579 | 5.87 | 1102 | 1.0787 |
| 38 | 0.26 | 71 | 0.003 | chr7:141441260-141441968 | 41 | 0.28 | 60 | 0.0024 |
| 20 | 0.15 | 21 | 0.003 | chr5:167945529-167946085 | 7 | 0.05 | 7 | 0.0008 |
| 2 | 0.01 | 2 | 0.000 | chr5:167945529-167946085 | 7 | 0.05 | 7 | 0.0008 |
| 284 | 2.96 | 587 | 0.498 | chr16:70065152-70065802 | 1097 | 11.44 | 3160 | 2.6833 |
| 150 | 1.59 | 189 | 0.121 | chr2:202335835-202337677 | 171 | 1.82 | 260 | 0.1671 |
| 30 | 0.23 | 39 | 0.005 | chr12:1004363-1005236 | 33 | 0.26 | 49 | 0.0061 |
| 1322 | 13.30 | 2407 | 1.877 | chr6:170061847-170062399 | 960 | 9.66 | 1533 | 1.1958 |
| 126 | 1.37 | 181 | 0.145 | Not Found | Not Found | 0.00 | Not Found | 0.0000 |
| 64 | 0.51 | 150 | 0.026 | chr3:107911374-107925474 | 68 | 0.54 | 157 | 0.0269 |
| 452 | 4.06 | 599 | 0.246 | chr4:3215464-3215684 | 738 | 6.62 | 1064 | 0.4371 |
| 373 | 3.59 | 605 | 0.203 | Not Found | Not Found | 0.00 | Not Found | 0.0000 |
| 86 | 1.28 | 120 | 0.161 | chr4:5743169-5743442 | 107 | 1.59 | 154 | 0.2071 |
| 29 | 0.27 | 51 | 0.034 | chr21:38794955-38844985 | 11 | 0.10 | 15 | 0.0099 |
| 23 | 0.23 | 28 | 0.012 | chr9:80535619-80537076 | 325 | 3.24 | 810 | 0.3614 |
| 312 | 2.67 | 1041 | 0.522 | chr1:35485881-35485983 | 413 | 3.53 | 1465 | 0.7348 |
| 191 | 1.38 | 286 | 0.025 | chr16:69482657-69492995 | 16 | 0.12 | 19 | 0.0017 |
| 41 | 0.46 | 58 | 0.064 | chr6:97649343-97676769 | 128 | 1.43 | 185 | 0.2029 |
| 167 | 1.92 | 230 | 0.232 | chr2:32112157-32117060 | 1010 | 11.64 | 1589 | 1.6014 |
| 1 | 0.01 | 1 | 0.000 | chr6:99868461-99873090 | 21 | 0.16 | 29 | 0.0033 |
| 1 | 0.02 | 1 | 0.001 | chr7:81705439-81713748 | 2 | 0.03 | 7 | 0.0073 |
| 38 | 0.26 | 71 | 0.003 | chr7:141441260-141441968 | 41 | 0.28 | 60 | 0.0024 |
| NA | NA | NA | NA | NA | NA | NA | NA | NA |
| 2429 | 21.22 | 16818 | 4.238 | chr8:131173040-131179781 | 2397 | 20.94 | 16510 | 4.1600 |
| Not Found | 0.00 | Not Found | 0.000 | chr14:19893151-19894699 | 31 | 4.86 | 34 | 2.2636 |
| 2728 | 24.06 | 6204 | 3.111 | chr7:32602526-32609631 | 811 | 7.15 | 1355 | 0.6796 |
| 29 | 0.27 | 51 | 0.034 | chr21:38794955-38844985 | 11 | 0.10 | 15 | 0.0099 |
| 351 | 3.71 | 481 | 0.488 | chrX:153741031-153741146 | 1834 | 19.39 | 3467 | 3.5144 |
| 210 | 2.89 | 295 | 0.260 | chr18:34322432-34322699 | 170 | 2.34 | 216 | 0.1906 |
| NA | NA | NA | NA | chr5:77070042-77072028 | 1306 | 9.21 | 4838 | 0.5132 |
| 51 | 0.41 | 103 | 0.020 | chr13:73299917-73301661 | 51 | 0.41 | 79 | 0.0153 |
| 20 | 8.37 | 22 | 6.984 | chr14:19682353-19683691 | 1 | 0.42 | 1 | 0.3175 |
| 233 | 1.80 | 467 | 0.053 | chr16:70578153-70578340 | 174 | 1.35 | 273 | 0.0310 |
| Not Found | 0.00 | Not Found | 0.000 | Not Found | Not Found | 0.00 | Not Found | 0.0000 |
| 124 | 1.10 | 195 | 0.116 | chr2:75929934-75933648 | 534 | 4.76 | 697 | 0.4146 |
| 778 | 6.12 | 1191 | 0.196 | chr17:36468625-36474585 | 847 | 6.66 | 1292 | 0.2123 |
| 414 | 3.86 | 568 | 0.358 | chr8:48186043-48192449 | 708 | 6.60 | 1131 | 0.7136 |
| 20 | 8.37 | 22 | 6.984 | chr14:19682353-19683691 | 1 | 0.42 | 1 | 0.3175 |
| 1472 | 11.38 | 3778 | 0.492 | chr16:15103419-15103537 | 116 | 0.90 | 190 | 0.0248 |
| 427 | 3.69 | 914 | 0.398 | chr1:117998829-118003110 | 40 | 0.35 | 57 | 0.0248 |
| 1841 | 14.37 | 4800 | 0.577 | chr3:12645037-12645634 | 2187 | 17.08 | 5789 | 0.6955 |
| 2778 | 62.81 | 7045 | 8.358 | chr20:34136962-34142142 | 194 | 4.39 | 922 | 1.0939 |
